# A novel mouse model of cerebral adrenoleukodystrophy highlights NLRP3 activity in lesion pathogenesis

**DOI:** 10.1101/2023.11.07.564025

**Authors:** Ezzat Hashemi, Isha Narain Srivastava, Alejandro Aguirre, Ezra Tilahan Yoseph, Esha Kaushal, Avni Awani, Jae Kyu. Ryu, Katerina Akassoglou, Shahrzad Talebian, Pauline Chu, Laura Pisani, Patricia Musolino, Lawrence Steinman, Kristian Doyle, William H Robinson, Orr Sharpe, Romain Cayrol, Paul Orchard, Troy Lund, Hannes Vogel, Max Lenail, May Htwe Han, Joshua Leith Bonkowsky, Keith P. Van Haren

## Abstract

**Objective:** We sought to create and characterize a mouse model of the inflammatory, cerebral demyelinating phenotype of X-linked adrenoleukodystrophy (ALD) that would facilitate the study of disease pathogenesis and therapy development. We also sought to cross-validate potential therapeutic targets such as fibrin, oxidative stress, and the NLRP3 inflammasome, in post-mortem human and murine brain tissues.

**Background:** ALD is caused by mutations in the gene *ABCD1* encoding a peroxisomal transporter. More than half of males with an *ABCD1* mutation develop the cerebral phenotype (cALD). Incomplete penetrance and absence of a genotype-phenotype correlation imply a role for environmental triggers. Mechanistic studies have been limited by the absence of a cALD phenotype in the *Abcd1-*null mouse.

**Methods:** We generated a cALD phenotype in 8-week-old, male *Abcd1-*null mice by deploying a two-hit method that combines cuprizone (CPZ) and experimental autoimmune encephalomyelitis (EAE) models. We employed *in vivo* MRI and post-mortem immunohistochemistry to evaluate myelin loss, astrogliosis, blood-brain barrier (BBB) disruption, immune cell infiltration, fibrin deposition, oxidative stress, and Nlrp3 inflammasome activation in mice. We used bead-based immunoassay and immunohistochemistry to evaluate IL-18 in CSF and post-mortem human cALD brain tissue.

**Results:** MRI studies revealed T2 hyperintensities and post-gadolinium enhancement in the medial corpus callosum of cALD mice, similar to human cALD lesions. Both human and mouse cALD lesions shared common histologic features of myelin phagocytosis, myelin loss, abundant microglial activation, T and B-cell infiltration, and astrogliosis. Compared to wild-type controls, *Abcd1*-null mice had more severe cerebral inflammation, demyelination, fibrin deposition, oxidative stress, and IL-18 activation. IL-18 immunoreactivity co-localized with macrophages/microglia in the perivascular region of both human and mouse brain tissue.

**Interpretation:** This novel mouse model of cALD suggests loss of *Abcd1* function predisposes to more severe cerebral inflammation, oxidative stress, fibrin deposition, and Nlrp3 pathway activation, which parallels the findings seen in humans with cALD. We expect this model to enable long-sought investigations into cALD mechanisms and accelerate development of candidate therapies for lesion prevention, cessation, and remyelination.

**Graphical Abstract:** 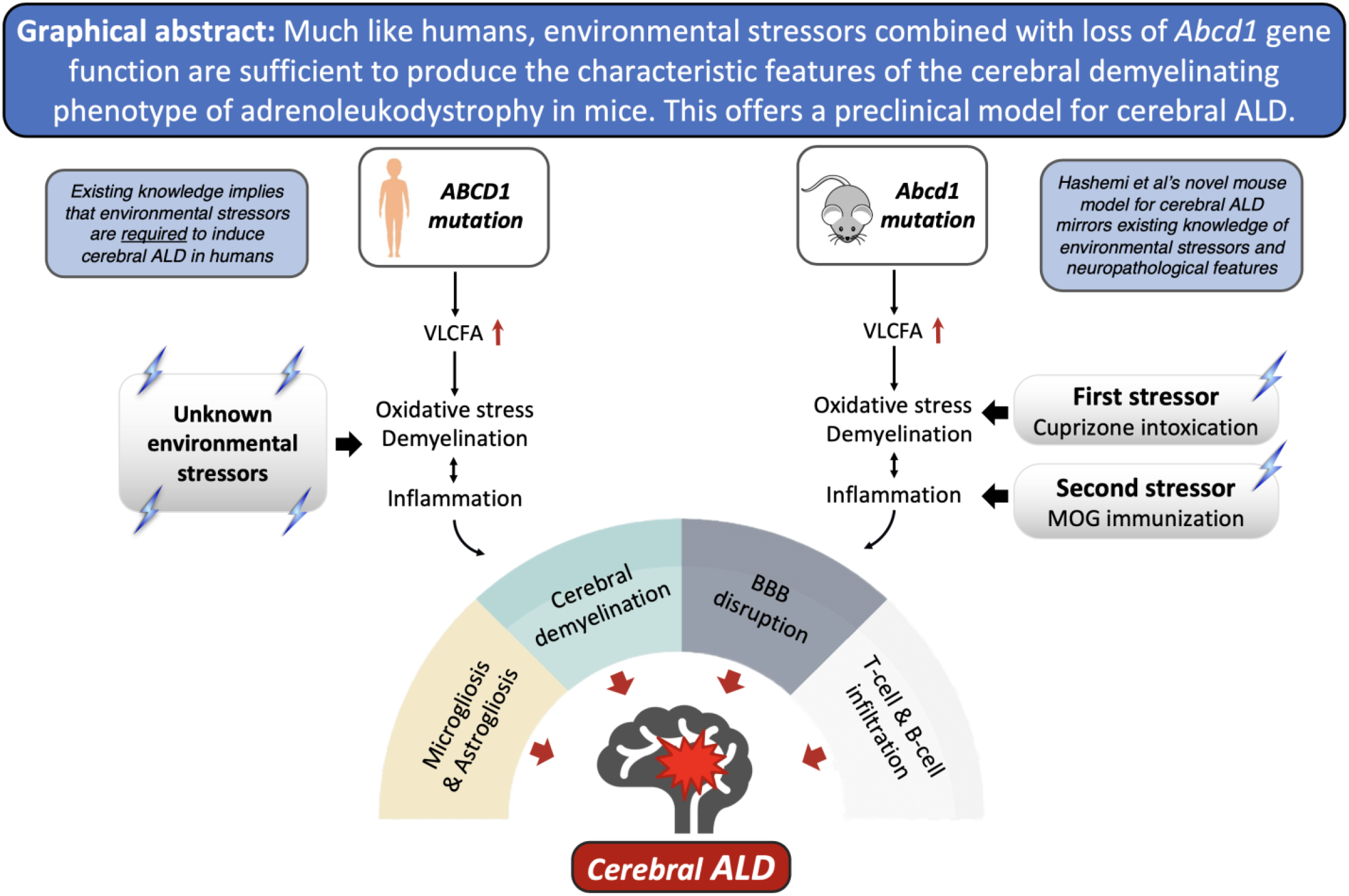

## Introduction

X-linked adrenoleukodystrophy (ALD) is an inherited neurodegenerative disease with a frequency of 1:15,000. It is caused by mutations in the *ABCD1* gene, which encodes a peroxisomal fatty acid transporter. ABCD1 mediates the transport of very long-chain fatty acids (VLCFAs) into the peroxisome for degradation. All affected males have elevated levels of VLCFAs in serum. Nearly all ALD males develop adrenal insufficiency and a progressive spastic paraplegia in a relatively predictable pattern by their fifth decade. In contrast, the incidence and timing of the cerebral demyelinating form of ALD, known as cerebral ALD (cALD), is more erratic. cALD has a bimodal risk distribution with the highest incidence in the first decade, where it affects roughly one-third of boys^1,2^. Although the risk of cALD rises again in the 3^rd^ and 4^th^ decades, approximately one-third of males with an *ABCD1* mutation never manifest cALD^3^.

cALD is a devastating, rapidly progressive form of inflammatory demyelination^4^. cALD lesions typically originate in the corpus callosum and are characterized by microglial activation, demyelination, BBB disruption, and infiltration of immune cells, primarily monocytes and T cells. The molecular pathogenesis of cALD lesions remains poorly understood. More recently, intraparenchymal fibrin deposition and, separately, NLRP3 activation have been implicated in cALD lesion pathogenesis^5,6^. Current effective treatments involve hematopoietic stem cell transplant with either donor cells or gene-corrected cells from the patient^7,8^. Unfortunately, transplant is limited to only a subset of children with early stage cALD, and associated with morbidity, mortality, complexity. Transplant has the potential to halt cALD progression but does not restore existing injury^9–11^. Therapy development for cALD has been hindered by a limited understanding of cALD pathophysiology and the absence of a preclinical model.

Currently, there is no preclinical model of cALD. Much like humans, *Abcd1*-null mice exhibit increased levels of VLCFAs in brain and immune tissue; they also develop signs and symptoms of myelopathy with advanced age^12^. However, *Abcd1*-null mice do not spontaneously manifest cerebral pathology^13^.

In both humans and mice, *Abcd1* deficiency alone is insufficient to induce cALD lesions. Although the trigger(s) for cALD in humans is unknown, oxidative stress and inflammation have been proposed as inciting events^14^. We hypothesized that a combination of oligodendrocyte oxidative stress (cuprizone diet, CPZ) with antigen-induced inflammatory demyelination (MOG-peptide EAE) would mimic human cALD brain lesions. To establish potential therapeutic targets in our mouse model, we sought to confirm the presence of NLRP3 inflammasome activation in human cALD tissue and evaluate the NLRP3 inflammasome, fibrinogen deposition and oxidative stress in our novel mouse model^5,6^.

We characterized the cerebral pathology of our novel mouse model by evaluating the following parameters: myelin loss, gliosis, immune cell infiltration (monocytes, T and B cells), perivascular cuffing, BBB breakdown, oxidative stress, and IL-18, a marker of NLRP3 inflammasome activation. To assess the viability of our model, we compared the histologic and molecular findings in our novel cALD mouse model to human cALD tissue as a gold standard and to similarly treated wild-type mice as a genotype control.

## Methods

### Post-mortem human tissue

Formalin-fixed brain tissue from three healthy pediatric controls and two pediatric patients with cerebral adrenoleukodystrophy (<21 years old) were provided by the NIH NeuroBioBank at the University of Maryland, Baltimore, MD. Tissue from healthy controls was age and gender-matched to patients with cALD. These specimens were obtained by consent at autopsy Gross pathology confirmed active demyelination in presumed brain lesions, validated through histological criteria^15,16^. Three active demyelination sites were analyzed per cALD patient.

### Cytokine assay and analysis

Cerebrospinal fluid (CSF) samples were collected from 20 cALD patients prior to hematopoietic stem cell transplantation at the University of Minnesota. Control CSF was collected from patients (n= 9) undergoing intrathecal chemotherapy as treatment for a prior diagnosis of acute lymphoblastic leukemia, who were at least 3 months into maintenance therapy and without CSF leukemia, as previously published^17^ . All subjects provided informed consent in accordance with the Declaration of Helsinki and were approved by the institutional review boards at the University of Minnesota and Stanford University. IL-18 levels were measured using ProcartaPlex Simplex bead sets (eBioscience) and a Bio-Plex instrument (Bio-Rad, Hercules, CA, USA). Median IL-18 concentrations and interquartile ranges were calculated for cALD and control groups, and significance was determined using a two-tailed Mann-Whitney U test.

### Experimental animals

C57Bl/6 wild-type and *Abcd1* knockout mice (Jax lab, B6.129-Abcd<tm1Kan>/J, Stock# 003716) were housed in groups of three to five with a 12-hour light/dark cycle at a controlled temperature with food and water provided ad libitum. The Institutional Animal Care and Use Committee at Stanford University approved all the animal protocols. In this study, animal protocols 32955 and 34033 were used and approved by the Administrative Panel on Laboratory Animal Care (APLAC) at Stanford University.

### Cuprizone intoxication and EAE induction

Eight-week-old *Abcd1^y/-^* and wild-type male mice were randomly assigned to one of four arms: no treatment (arm 1), cuprizone (CPZ) treatment alone (arm 2), EAE treatment alone (arm 3) or combination of CPZ/EAE treatment (arm 4). A total of 57 mice (n=29 *Abcd1^y/-^* and n=28 wild-type) were used in the study. **Supplemental Table S1** details the distribution of mice across each experimental arm and assay.

For CPZ treatment alone, mice were initially on a standard chow diet for two weeks, followed by a 2-week 0.2% CPZ diet (TD.140800, ENVIGO) as previously described^18^. This duration of CPZ treatment was chosen for its significant immune cell infiltration and loss of mature oligodendrocytes in the brain (**Supplemental Fig 1A**)^18^.

For EAE alone, mice were immunized with 100 µg of MOG_35-55_ (Genemed synthesis, Inc) in an emulsion containing complete Freund’s adjuvant (CFA, Difco), then injected peritoneally with 200 ng of Bordetella pertussis toxin (lot# 181236A1, List Biological Laboratories) on post-immunization day 0 and 2. For CPZ/EAE treatment, *Abcd1^y/-^* and wild-type mice were fed 0.2% CPZ diet for 2 weeks followed by MOG immunization^18^. We performed two separate rounds of CPZ alone and EAE alone treatments, and we also conducted CPZ/EAE combined treatment at three different time points. We used 2-3 mice per group in each round.

The experimental design was structured to collect samples during the peak of disability between days 16 and 22 following the injection. In arms 3 and 4, we monitored mice and scored EAE clinical symptoms after immunization based on previous study^19^. We collected tissue samples during this timeframe when the mice reached disability scores of 2 or 3. We also collected the samples on day 22 if the mice displayed no clinical sign of EAE or had score below 2. Therefore, the motor disability scores are demonstrated till day 16 to include all mice in the clinical scoring analysis. No mice were excluded from analysis.

### Mouse in vivo MRI acquisition and processing

We obtained *in vivo* MRI images utilizing an actively shielded Bruker horizontal bore scanner (Bruker Corp, Billerica MA) with 11.7 T field strength and International Electric Co. (IECO) 750/400 gradient drivers at the Stanford Center for Innovation in *In vivo* Imaging (SCi3). For each mouse, we obtained T1 and T2-weighted images. Gadolinium (Gadavist) at a 100 mg/kg dose was administered using an intraperitoneal injection 30 minutes prior to imaging^20^. We used OsiriX software to quantify intensity signaling and analyze myelin loss in approximately 0.7 mm^2^ of the medial corpus callosum (MCC) in the rostral diencephalon.

### Immunofluorescence and immunohistochemistry

The collection of tissue, luxol fast blue (LFB) staining, and immunofluorescence were performed based on the procedure detailed in our previous study ^18^. The **Supplemental Table S2** contains the details of the antibodies including their concentrations and manufacturing information.

### Quantification based on grading and intensity

All experiments were assessed by a trained reader blinded to the identity and treatment of the mouse. Cell counts were conducted in 0.1 mm^2^ of MCC and normalized to DAPI. ImageJ was used for cell counting and intensity measurements. Final counts/scores were averaged from 2-3 tissue sections of tissue per mouse. Brain tissue analysis was focused between the Bregma −2.355 and −1.255 mm in mice.

To evaluate the severity of perivascular cuffs (PVCs), we quantified the PVCs in 2 to 3 brain sections of each mouse. We used DAPI staining to analyze the degree of perivascular immune cell infiltration and its severity was graded on the scale of I to IV as follows: Grade I: leukocyte foci with an aggregation of inflammatory cells; Grade II: accumulation of immune cells around the vessels without parenchymal penetration; Grade III: penetration of immune cells into the parenchyma, and Grade IV: prominent immune cell penetration and creation of a large lesion (**Supplemental Fig 2A**). The same grading scheme was applied for evaluating CD68^+^ macrophage/microglia infiltration (**Supplemental Fig 2B**). Perivascular infiltration of T and B cells was graded similarly, with one exception that Grade I represented fewer cells attached to endothelial cells in the vessels (**Supplemental Fig 2C**).

We graded astrogliosis on a grade of I - IV based on the intensity of GFAP staining in the MCC (**Supplemental Fig 2D**), whereby grade I represented minimal or no astrogliosis and grade IV represents severe lesions. The severity of demyelination in LFB staining was graded based on the intensity of LFB in MCC. (**Supplemental Fig 3**).

### Statistical analyses

We analyzed the data with GraphPad Prism 9 and tested the normality of distribution by using the Shapiro-Wilk test. Significant differences between two groups were determined using unpaired t-tests with Welch’s correction. For multiple group comparisons, one-way ANOVA was utilized with Brown-Forsythe for variance homogeneity in normally distributed groups and Kruskal-Wallis for non-normal distributions. False discovery rate (FDR) correction by Benjamini and Hochberg method ensured accurate multiple comparisons (q < 0.05). Results were presented as mean ± SD in graphs. Sample size estimation using G*Power 3.1 recommended 5 mice per group for 90% statistical power, based on effect size estimated from previous MBP staining data^18^.

## Results

### cALD donors had advanced cerebral disease at time of diagnosis

**Figure 1** highlights key radiologic and histologic features from two males affected by cALD. cALD donor 1 was 7 years old at the time of autopsy. Brain MRI, obtained 10 months prior to death, demonstrates extensive diffuse contiguous symmetric T2 hyperintense lesions involving frontotemporal, central, and anterosuperior occipitoparietal white matter. These lesions exhibited gadolinium contrast enhancement and received MRI severity score (Loes score) of 16 as depicted in **Fig 1A**. His clinical course was complicated by primary adrenal insufficiency, seizures, g-tube dependence and loss of consistent bowel and bladder though he continued to ambulate independently.

**Figure 1.**
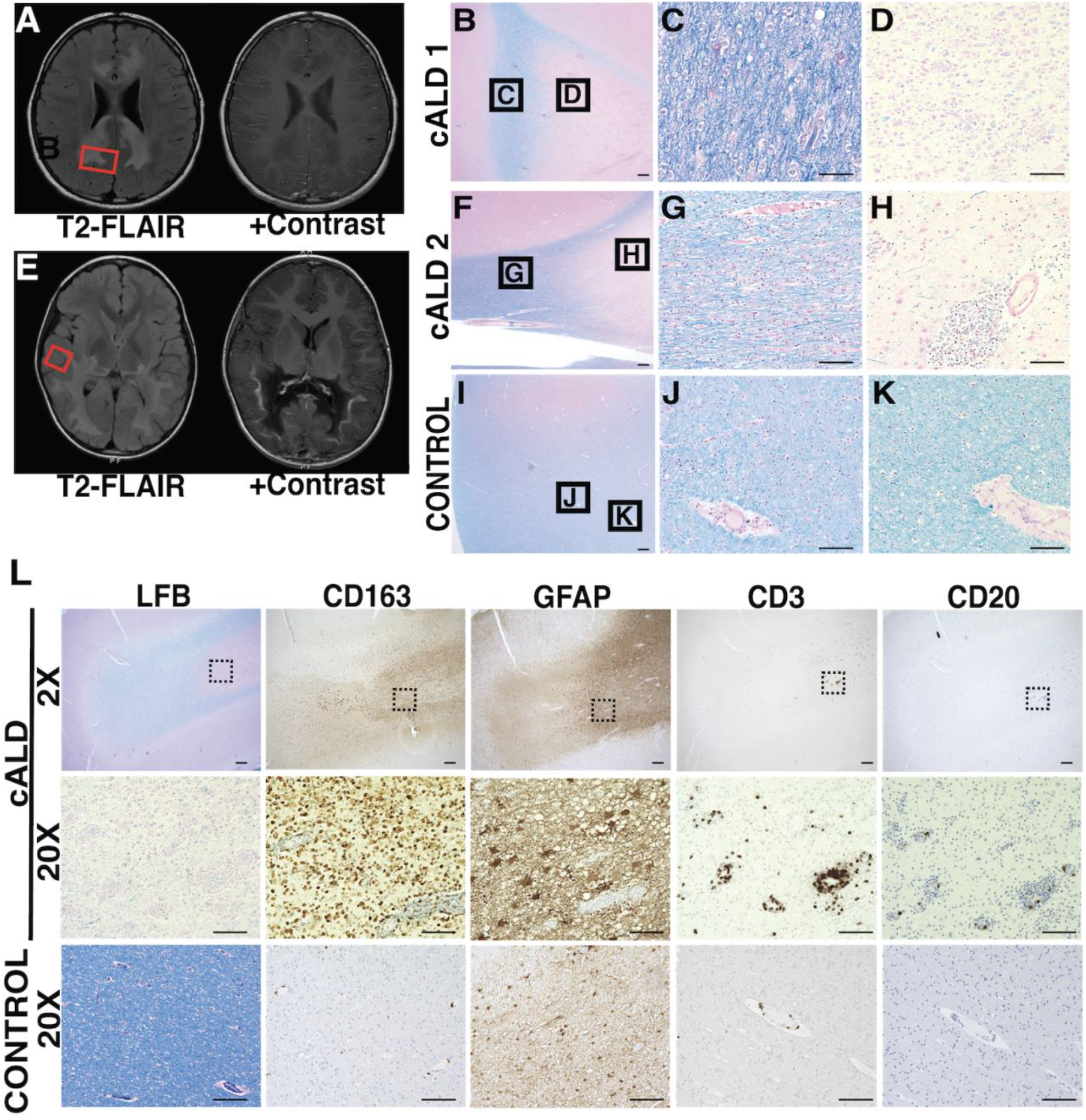
In boys with cALD, brain lesions exhibit important radiologic and histologic hallmarks. (A, E) Axial MR imaging from two children with cALD highlight areas of demyelination (T2 FLAIR intensities) and vascular breakdown (contrast enhancement). Red boxes indicate representative areas sampled for histopathological images. (B-K) LFB + H&E histology of postmortem tissue from cALD patients with varying severity of demyelination in lesions. Histology highlights regions of extensive, active demyelination (C, G) and early gliosis (D, H) compared to healthy, unaffected control (I-K). B, F, I: 2X magnification of representative sections, scale bar: 400um. C, D, G, H, J, K: 20X magnificent of respective sections, scale bar: 100um. (L) Immunohistochemistry of representative demyelinated cALD lesion shows infiltration of microglia/ macrophages (CD163), and to a lesser extent T-cells (CD3) within the lesion. B cells (CD20) are present in lower abundance. Scale bar for 2X magnification represents 400µm; black boxes indicate corresponding areas shown at 20X magnification. Scale bar for 20X magnification represents 100µm

cALD donor 2 was 10 years old at the time of autopsy. Prior to his diagnosis, he had experienced progressive behavioral changes and vision processing disorder. Brain MRI, obtained 28 months prior to death, demonstrated extensive T2 hyperintense, posterior predominant, confluent lesions including involvement of splenium with contrast enhancement, and received a Loes score of 13 (**Fig 1E**). His clinical course was complicated by adrenal insufficiency.

Both donors had *ABCD1* mutations and high plasma VLCFA levels. Given their extensive brain lesions at diagnosis, neither donor was eligible for transplant therapy.

### Perivascular immune cell infiltration in human cALD

cALD brain lesions radiate outward creating distinct concentric histopathological regions with a central area of gliosis (**Fig 1D, H**), surrounded by concentric areas of progressive demyelination (**Fig 1C, G**). We confirmed that the human brain tissue sections had perilesional, active demyelination, and gliosis regions based on previously defined criteria^15,16^. Histology and LFB staining identified active demyelination areas and extensive inflammatory infiltration of macrophages/microglia, T and B cells (**Fig 1L**). Sections also contained regions consistent with perilesional white matter characterized by pale LFB staining and scattered macrophage/microglia immunoreactivity (**Fig 1B, D, F, H**).

We noted perivascular infiltration consisting predominantly of macrophages/ microglia, with a distinct absence of astrocytes in the immediate perivascular area (**Fig 1L**). Furthermore, there was a consistent presence of CD3^+^ T-cells in the perivascular region (**Fig 1L**). In agreement with prior literature^21^, perivascular CD20^+^ B-cell infiltration was inconsistent and present to a lesser degree compared to T-cells, and macrophages/microglia (**Fig 1L**).

### CSF and brain tissue from boys with cALD show evidence of NLRP3 inflammasome activation

IL-18 levels in CSF were elevated in cALD compared to controls (median ± IQR), 12.89 (10.04 −22.83) and 5.56 (5.56-6.22) respectively (**Fig 2A**). IL-18 immunoreactive cells resembled the morphology and distribution of microglia/macrophages, and astrocytes (**Fig 2B, C**). Perivascular IL-18 co-localized exclusively with Iba1^+^ microglia/macrophages (**Fig 2C**).

**Figure 2.**
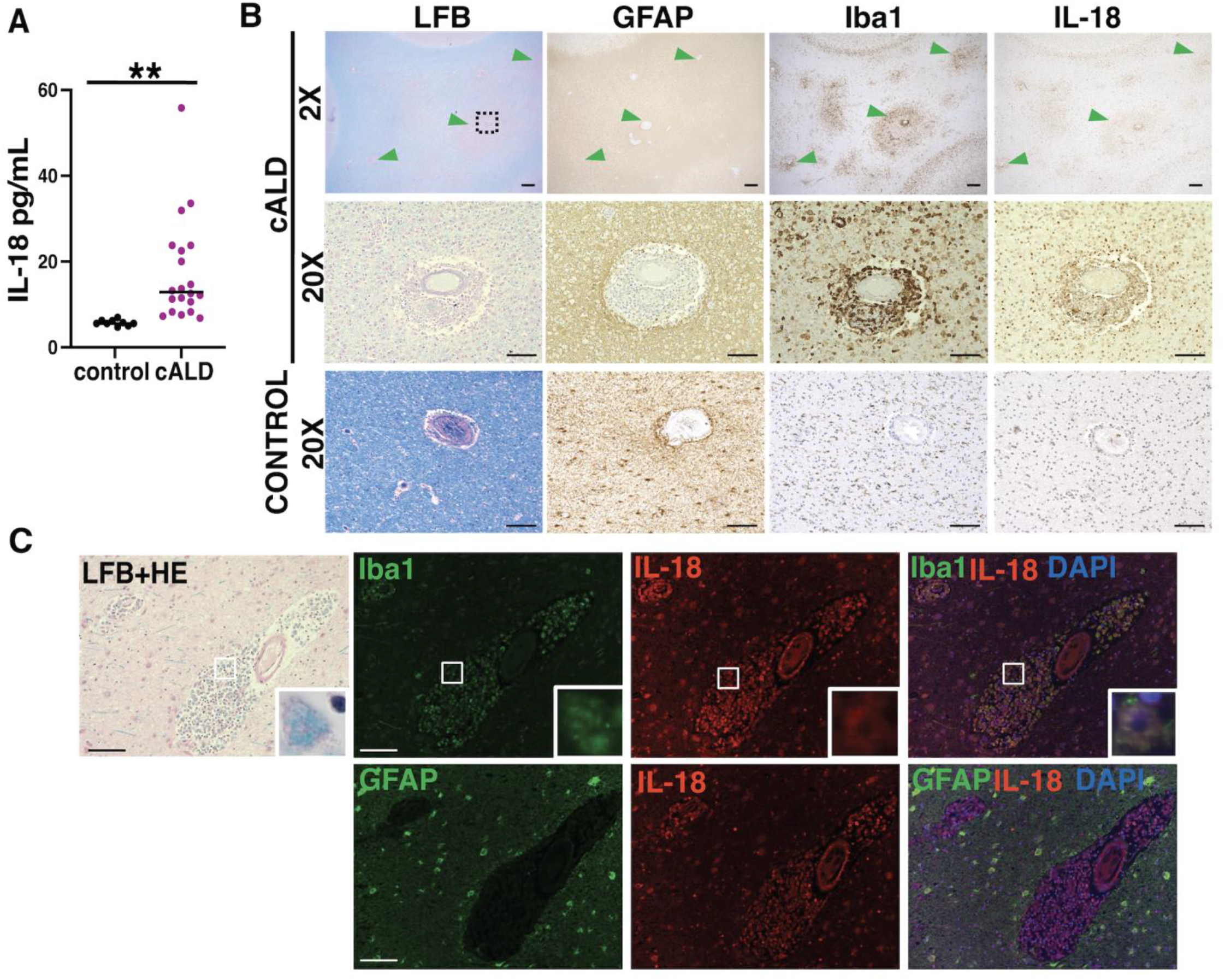
In boys with cALD, interleukin-18, an NLRP3 inflammasome effector molecule, is increased in spinal fluid and brain tissue. (A) IL-18 levels are significantly higher in the CSF of children with cALD compared to unaffected, age-matched controls (**: p < 0.001). (B) Within areas of early brain demyelination in cALD (green arrowheads), the distribution of IL-18^+^ cells align with microglia (Iba1) rather than astrocytes (GFAP). Black boxes indicate corresponding areas shown at 20X magnification. (C) Immunofluorescence highlights co-localization of IL-18 with perivascular microglia/macrophages (Iba1^+^) and, to a lesser extent, intraparenchymal astrocytes (GFAP). Insets highlight co-localization at 60X magnification. 2X magnification scale bar: 400µm. 20X magnification scale bar: 100µm.

### Combined CPZ/EAE induces a robust neurologic phenotype in Abcd1^y/-^ mice

We hypothesized that the combination of *Abcd1* dysfunction with CPZ/EAE leads to cerebral inflammation characteristic of human cALD. To test this, we used CPZ and EAE alone as controls and in combination in 8-week-old *Abcd1^y/-^* and wild-type mice (**Fig 3A**). The motor disability scores for all experimental arms are presented in **Fig 3B**. *Abcd1^y/-^* mice with CPZ/EAE treatment (cALD mice) exhibited a more severe phenotype compared to wild-type mice on post-immunization days 12 and 13 (*p*=0.04, and *p*=0.03, respectively) (**Fig 3C**). Compared to EAE alone, the cALD mice showed a more severe neurologic phenotype (motor disability in tail and/or limbs) post-immunization day 11, 12 13 (with *p*=0.02, *p*=0.001, and *p*=0.03, respectively in **Fig 3D**), and greater aggregate motor disability scores *p*=0.03 (**Fig 3E**).

**Figure 3.**
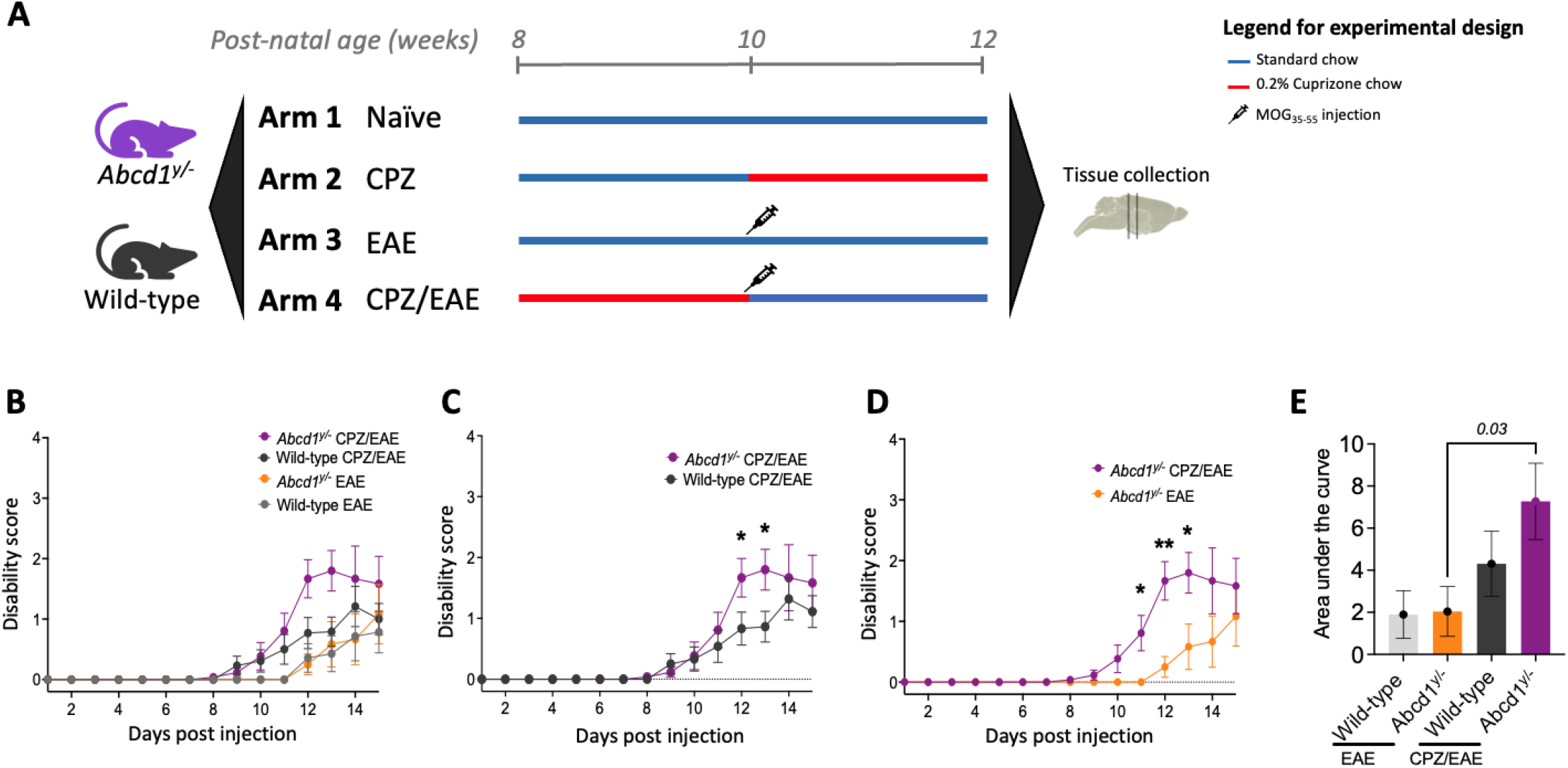
In *Abcd1*-null mice, exposure to a combination of CPZ/EAE induced greater motor disability than similarly treated wild-type controls or treatment with CPZ or EAE alone. (A) A graphical overview summarizes the treatment regimens and timelines for the four experimental arms applied to both *Abcd1*-null and wild-type C57BL/6J mice. (B) Clinical disability scores were higher among mice in CPZ/EAE arm (n=12-13/genotype) compared with EAE (n=6-7/genotype); mice in the CPZ arm (n=6) had no measurable disability. (C) Clinical scores in wild-type and *Abcd1*y/-mice in CPZ/EAE models (n=12-13 male mice in each group). (D, E) *Abcd1*^y/-^ mice induced with CPZ/EAE had significantly higher disability than mice treated with EAE alone. The bar graph represents the mean ± SEM of motor disability scores. The unpaired t-test determined the significant difference between the two groups; p*<0.05 was considered significant. CPZ: Cuprizone; EAE: experimental autoimmune encephalomyelitis; ROI: region of interest

In EAE alone, 7 out of 7 wild-type and 3 out of 6 *Abcd1^y/-^*mice developed motor disability. However, in CPZ/EAE, 11 out of 13 wild-type mice and 12 out of 13 *Abcd1^y/-^* mice developed motor disability. Both wild-type and *Abcd1^y/-^*mice treated with CPZ/EAE exhibited signs of inflammatory disease progression beginning around day 10 after the immunization.

In the CPZ/EAE treatment group, wild-type mice showed no significant difference in neurologic phenotype compared to EAE alone-(**Supplemental Fig 3A).** In EAE alone, *Abcd1^y/-^* deficiency had no significant impact on the EAE clinical scores (**Supplemental Fig 3B**). There was no difference in disease onset timing between wild-type and *Abcd1^y/-^* mice in the CPZ/EAE model (**Supplemental Fig 3C**).

### Cuprizone/EAE in Abcd1^y/-^ mice induces cerebral demyelination, BBB disruption, and oxidative stress similar to human cALD

We examined whether the cALD mouse model developed cerebral demyelination, a hallmark of human cALD pathology. We focused on characterizing demyelinating in the MCC, given that human cALD brain lesions are posterior predominant and begin medially in the splenium^22,23^. As with human cALD, T2 hyperintensity in the cALD was most pronounced in the MCC and significantly higher than both CPZ/EAE-treated wild-type mice and *Abcd1^y/-^* naïve mice, *p*=0.007 and *p*=0.003, respectively (**Fig 4A, B**).

**Figure 4.**
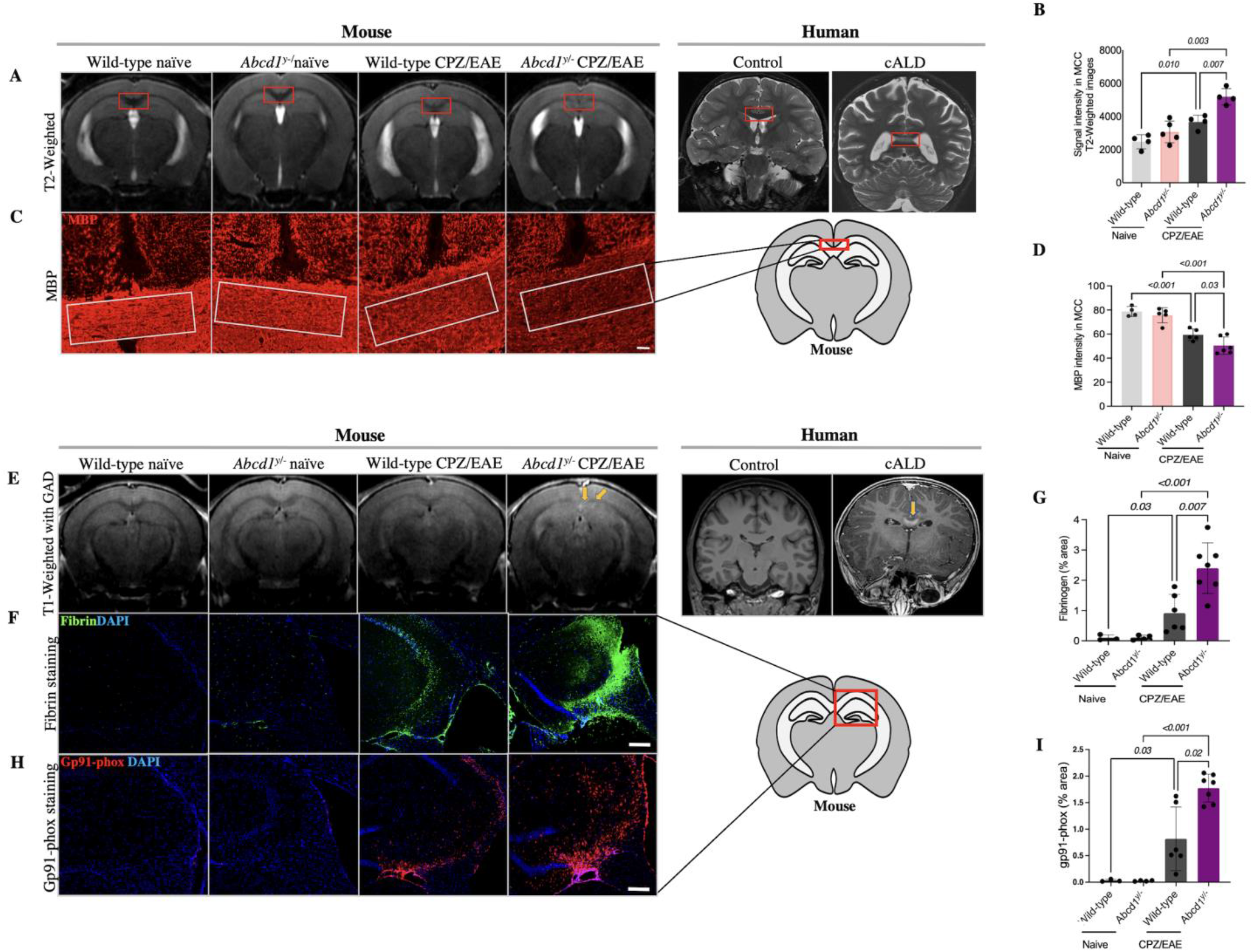
*Abcd1*-null mice induced with CPZ/EAE exhibit radiologic and histological features of demyelination and blood brain barrier disruption. (A) Representative coronal MRI images demonstrate high T2 signal in the MCC (red square) of *Abcd1^y/-^* induced with CPZ/EAE resembling. the characteristic pattern seen in human cALD MRI (far right). (B) A comparison of quantitative analyses of T2-weighted MRI of the MCC shows a higher T2 signal in the MCC (highlighted by red box) of *Abcd1*^y/-^ mice induced with CPZ/EAE, similar to early lesions in human cALD. (C) Representative histologic sections of MBP staining in the MCC depict differing levels of demyelination across experimental arms. White box denotes representative areas of quantitative analysis. Scale bars, 50µm. (D) Quantitative analysis of MBP intensity in the MCC shows high levels of demyelination in *Abcd1*^y/-^ mice induced with CPZ/EAE. (E) Representative images of post-gadolinium T1-weighted MRI in show focal areas of gadolinium enhancement (yellow arrows) in *Abcd1*^y/-^ mice treated with CPZ/EAE, similar to characteristic findings in human cALD. (F, G) Representative images and quantitative measures of immunostaining for fibrinogen, a marker of BBB disruption, show high levels in *Abcd1*^y/-^ mice induced with CPZ/EAE. (H, I) Similarly, representative and quantitative analysis of immunostaining for gp91-phox, a marker of oxidative stress, are highest a in *Abcd1*^y/-^ mice induced with CPZ/EAE. Scale bars, 200 µm. All bar graphs display the mean ± SD. Statistical comparisons via one-way. MCC: medial corpus callosum; MBP: myelin basic protein; CPZ: cuprizone; EAE: experimental autoimmune encephalomyelitis.

MBP staining revealed greater myelin loss in the MCC of cALD mice compared to CPZ/EAE-treated wild-type mice and *Abcd1^y/-^* naïve mice (*p*= 0.03 and *p*<0.001, respectively) (**Fig 4C, D**). LFB staining further confirmed myeline loss in *Abcd1^y/-^* mice compared to wild-type mice in the CPZ/EAE treatment group (**Supplemental Fig 4A, B**). Due to the extensive demyelination, we quantified oligodendrocytes in the MCC. Our analysis revealed a significant decrease in Olig2^+^ cells within MCC in mice with CPZ/EAE treatment compared to naïve mice, *p*=0.01 (**Supplemental Fig 5A, B**).

Human cALD is characterized by BBB disruption and elevated levels of oxidative stress^24,25^. To assess BBB integrity, we used gadolinium and T1 weighted MRI sequences obtained 5 weeks after initiating CPZ/EAE. We observed gadolinium enhancement at the MCC lesion site, resembling enhancement seen in human cALD (**Fig 4E**). We used immunostaining to quantify deposition of blood-derived fibrin in the brain as a measure of BBB disruption. cALD mice showed higher fibrin deposition compared to wild-type mice in CPZ/EAE model, *p*=0.007 (**Fig 4F, G**). To evaluate oxidative stress, we used immunostaining for oxidative stress marker NADPH oxidase subunit, gp91-phox, which is upregulated in cALD mice compared to wild-type mice in CPZ/EAE model, *p*=0.02 (**Fig 4H, I**). **Supplemental Table S3** provides mean ± SD for each group in MRI and immunofluorescence tests.

### cALD mice exhibit perivascular infiltration of macrophages/microglia and lymphocytes

PVCs consist of immune cell infiltration in the perivascular space and have been documented in the brain following viral or immune-mediated inflammatory processes^26,27^. **Fig 5A** demonstrates the spatial distribution of PVCs in cALD mice. To analyze PVC severity, we employed a grading system based on infiltration depth and lesion size. *Abcd1^y/-^* and wild-type mice in CPZ or EAE alone groups had few PVCs, and the PVCs were low grade (**Fig 5A**). The cALD mouse model showed significantly higher grade of PVCs both compared to wild-type mice in CPZ/EAE group, *p*=0.01 (**Fig 5D**), as well as *Abcd1^y/-^* in CPZ or EAE alone groups *p*=0.001 (**Fig 5D**). These findings replicate the perivascular infiltration seen in cALD human pathology (**Fig 2B**).

**Figure 5.**
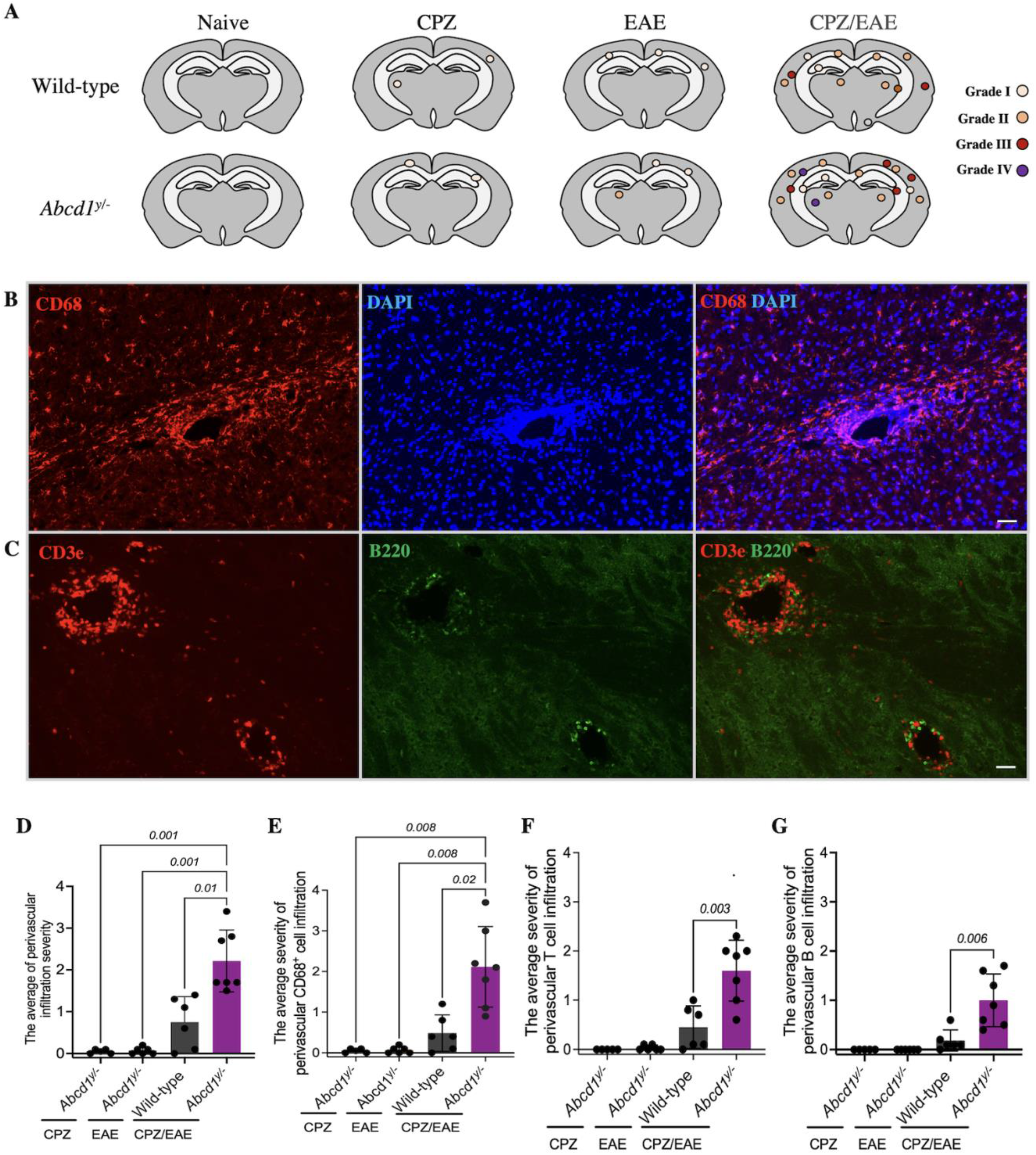
*Abcd1*-null mice induced with CPZ/EAE exhibit abundant perivascular infiltration of macrophages, T-cells, and B-cells. (A) The cartoons illustrate the spatial distribution of perivascular cuffs in a representative mouse, as determined by DAPI staining, across experimental arms. Representative histologic images depict perivascular infiltration of (B) macrophage/microglia (CD68 staining) (C) T cells (CD3e staining) and B cells (B220 staining) in the *Abcd1*^y/-^ mice induced with CPZ/EAE similar to human cALD. Scale bars, 50 µm. Quantitative assessment shows high levels of (D) total immune cells, (E) macrophages/microglia, (F) T cells, and (G) B cells in *Abcd1*^y/-^ mice induced with CPZ/EAE. Bar graphs display the mean ± SD. Brown-Forsythe Welch ANOVA test determined the significance between the groups. CPZ: cuprizone; EAE: experimental autoimmune encephalomyelitis.

We further analyzed the cellular composition of perivascular infiltration by evaluating macrophages/microglia, T-cells, and B-cells using immunohistochemistry. The severity of macrophages/microglia perivascular infiltration is higher in cALD mice compared to wild-type in CPZ/EAE group, *p*= 0.02 (**Fig 5B, E**). The severity of T cell (**Fig 5C, F**) and B cell (**Fig 5C, G**) perivascular infiltration is higher in cALD mouse model compared to wild-type in CPZ/EAE model, *p*= 0.003 and *p*= 0.006, respectively.

### cALD mice exhibit microgliosis and astrogliosis

To investigate activated macrophages/microglia in our new cALD mouse model, we conducted CD68^+^ immunostaining. CPZ treatment alone increases the CD68^+^ cells in the MCC of both *Abcd1^y/-^* and wild-type mice. In the CPZ/EAE treatment, *Abcd1^y/-^* and wild-type mice displayed higher CD68^+^ cells in the MCC than in EAE treatment alone, *p*<0.001 and *p*=0.03, respectively. However, there was no statistically significant difference between wild-type and *Abcd1^y/-^* in CPZ/EAE, EAE, and CPZ models. CD68^+^ cells in *Abcd1 ^y/-^* and wild-type mice in CPZ/EAE model are increased compared to CPZ alone but did not reach statistical significance (**Fig 6A, C**).

**Figure 6.**
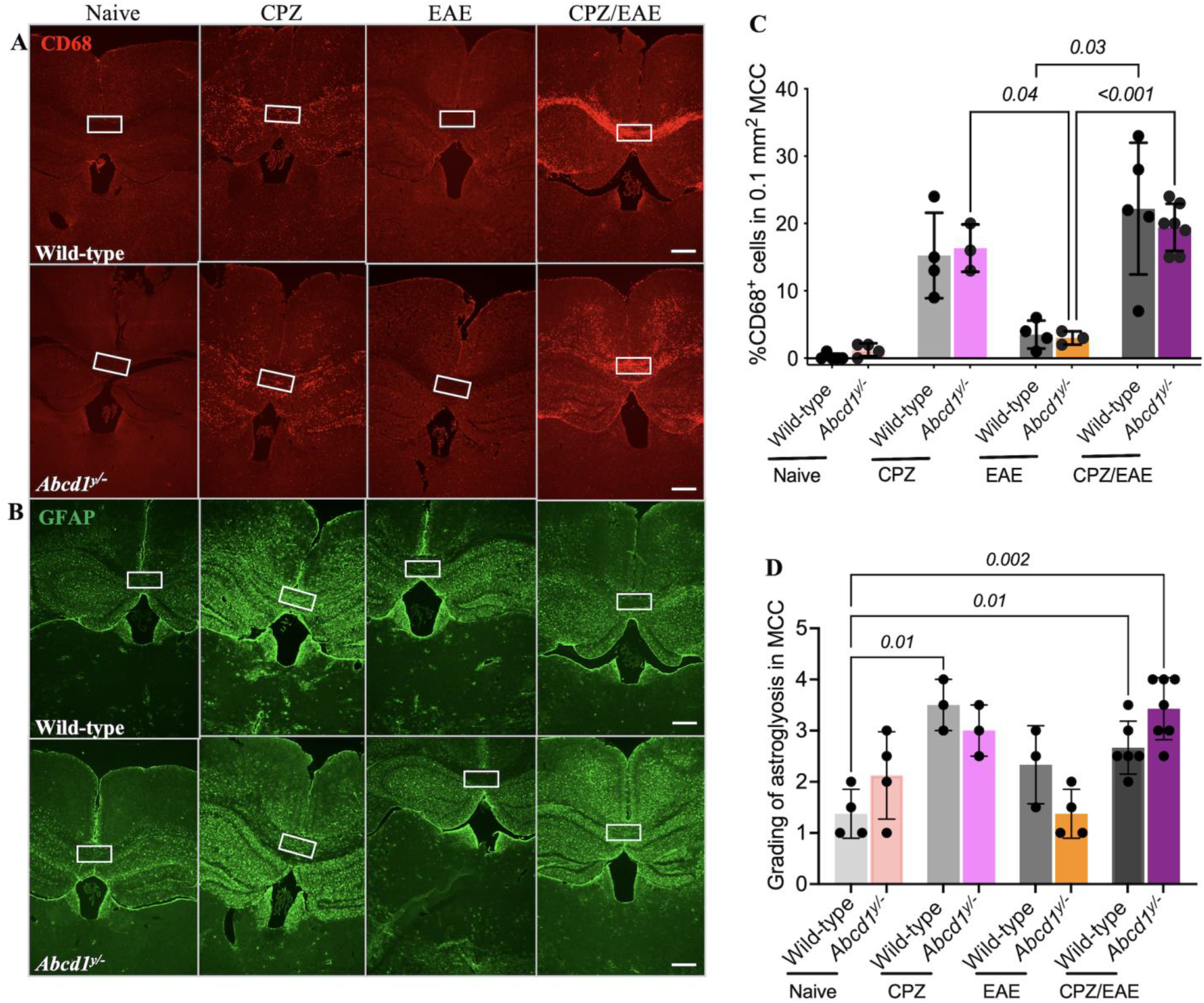
*Abcd1*-null mice induced with CPZ/EAE exhibit increased microgliosis and astrogliosis in the cerebral white matter. Representative histologic sections show (A) CD68 and (B) GFAP staining across the four experimental arms. The white square highlights the MCC. Quantitative analysis of the MCC shows elevated (C) CD68^+^ cell count and (D) GFAP grading in the CPZ and CPZ/EAE experimental arms. Bar graphs display the mean ± SD. Statistics calculated via one-way ANOVA. Scale bar: 400µm. CPZ: cuprizone; EAE: experimental autoimmune encephalomyelitis; MCC: medial corpus callosum.

We examined astrogliosis using GFAP staining. In the MCC, there was no significant difference in astrogliosis between *Abcd1^y/-^* and wild-type mice in the naïve group (**Fig 6B**). CD68 and GFAP staining in MCC at higher magnification is shown in **Supplemental Fig 6A, B**. CPZ treatment alone increased astrogliosis in wild-type mice, *p*=0.01 (**Fig 6D**). In the CPZ/EAE group, *Abcd1^y/-^* mice had higher astrogliosis than wild-type mice, but it was statistically insignificant. These findings indicate no significant difference in astrocytes and CD68^+^ cells in the MCC of wild-type and *Abcd1^y/-^* mice after CPZ/EAE treatment.

### cALD mice exhibit perivascular demyelination and myelin phagocytosis

In human cALD, affected white matter contains perivascular inflammatory cells and demyelination. Our mouse animal model revealed similar perivascular demyelination and accumulation of CD68^+^ cells around the vessels (**Fig 7A**). Perivascular macrophages/microglia indicate myelin inside the cells, which might be due to myelin phagocytosis (**Fig 7B**).

**Figure 7.**
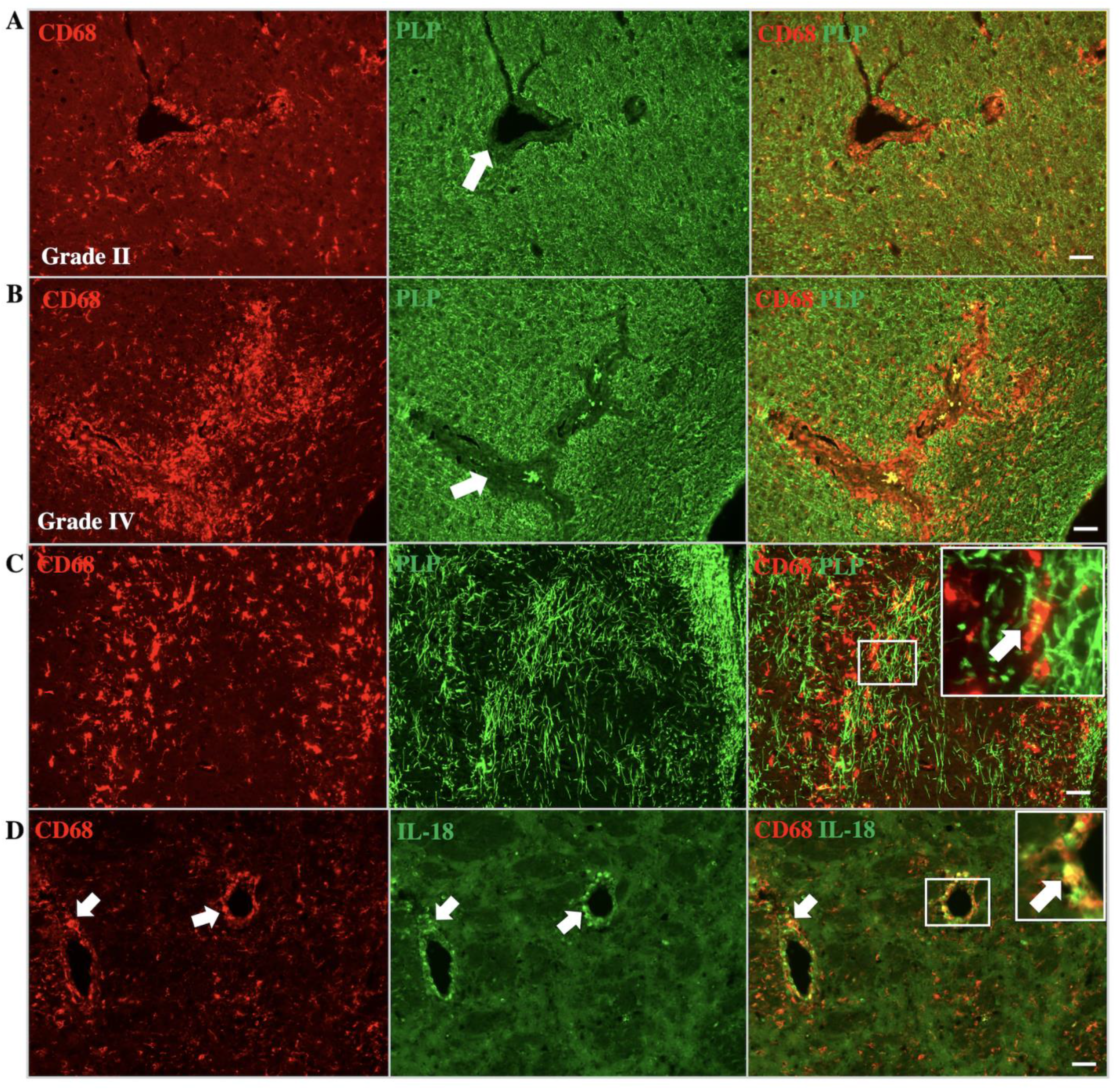
*Abcd1*-null mice induced with CPZ/EAE show perivascular demyelination, myelin phagocytosis, and IL-18 expression in macrophages/microglia. Co-staining with CD68 (macrophage/microglia) and PLP (myelin) indicate macrophage/microglia infiltration and demyelination in perivascular white matter. The images represent (**A**) grade II and (**B**) grade IV perivascular infiltration of macrophage/microglia with associated demyelination. White arrows indicate perivascular regions with associated loss of myelin. (**C**) The images display phagocytosis of myelin by macrophage/microglia. Inset depicts higher magnification of myelin phagocytosis in the corresponding region. (**D**) IL-18 expression in perivascular macrophage/microglia, similar to human cALD. Arrows indicate colocalization. Inset depicts a higher magnification of IL18 and CD68 colocalization in the corresponding region. Scale bar: 50µm. CPZ: cuprizone, EAE; experimental autoimmune encephalomyelitis; PLP: proteolipid protein.

### cALD mice exhibit IL-18 expression in intraparenchymal macrophages and microglia

Our findings of human postmortem cALD brain tissue indicates increased IL-18 immunoreactivity in perivascular macrophages/microglia. The cALD mouse model also demonstrates IL-18 immunoreactivity in perivascular macrophages/microglia (**Fig 7C**). We also observed colocalization of CD68 and IL-18 in the lateral corpus callosum of *Abcd1^y/-^* and wild-type mice in CPZ/EAE model, which is higher in *Abcd1^y/-^* mice (**Supplemental** Fig 7).

### cALD mice exhibit sustained neuroinflammation and cerebral demyelination

In cALD mice, demyelination persists in the MCC after 10 weeks of initiating CPZ/EAE induction, confirmed by T2-weighted MRI (n=3-5) (**Supplemental Fig 8A**). Additionally, the mice exhibited sustained microgliosis in the MCC after 10 weeks of starting the CPZ/EAE induction, with a statistically significant difference (P=0.05) compared to the naïve group. However, we did not observe astrogliosis in the MCC at this time point (**Supplemental Fig 8B, C)**.

## DISCUSSION

More than half of boys and men with a mutation in *ABCD1* will develop inflammatory cerebral demyelination (cALD), which can be fatal if untreated and remains poorly understood. The absence of spontaneous cerebral demyelination in *Abcd1*-null mice has hampered investigations into both the mechanism and treatment of cALD for decades. We present new evidence supporting NLRP3 activation as a driver of neuroinflammation in human cALD tissue^6,28^ . We also describe a two-hit induction method combining CPZ and MOG-EAE that induces a robust cerebral demyelinating phenotype in male *Abcd1*-null mice and recapitulates key radiologic, histologic, and molecular features of cALD, including NLRP3 inflammasome activation, BBB disruption, fibrin deposition, and oxidative stress (**Table 1**). Collectively, these findings suggest that our mouse model offers new avenues for investigating both disease mechanisms and therapy development for cALD.

**Table 1:** The characteristic features of the cerebral demyelinating phenotype in boys and men with *ABCD1* mutations can be reproduced in *Abcd1*-null mice following exposure to a combination of CPZ diet and MOG-peptide EAE injections.

| Characteristic features of brain lesions in humans with the cerebral ALD phenotype | Features observed in mice with *Abcd1*<sup>y/-</sup> genotype |  |  |  |
| --- | --- | --- | --- | --- |
|  | No induction | EAE alone | CPZ alone | CPZ + EAE |
| Elevated VLCFAs in brain and immune cells | + | + | + | + |
| Microglial activation in the cerebral white matter | - | - | + | + |
| Macrophage-mediated phagocytosis of cerebral myelin | - | - | + | + |
| Cerebral demyelination centered over the corpus callosum | - | - | + | + |
| Elevated markers of oxidative stress | - | - | + | + |
| NLRP3/IL-18 pathway activation | - | - | + | + |
| Cerebral astrogliosis | - | - | + | + |
| Blood-brain barrier disruption with fibrin deposition | - | - | - | + |
| Perivascular aggregation of macrophages, T-cells, B-cells | - | - | - | + |
| Infiltration of T-cells into brain parenchyma | - | - | - | + |
| Infiltration of B-cells into brain parenchyma | - | - | - | + |
| Perivascular demyelination | - | - | - | + |

*Abbreviations: ALD: adrenoleukodystrophy; CPZ: cuprizone; EAE: experimental autoimmune encephalitis; VLCFAs: very long chain fatty acids*

As with other demyelinating disorders, the triggers for cALD are largely unknown, but both oxidative stressors and inflammatory triggers have been proposed^29,30^. These notions underpinned our rationale for combining CPZ, an oxidative stressor, and MOG-EAE, an antigenic trigger, to induce a cALD phenotype in *Abcd1*-null mice. Our model complements the efforts of investigators who have recently begun combining CPZ and EAE in wild-type mice to overcome some of the limitations of stand-alone EAE and CPZ models for demyelination inflammation^31–33^. Traditional EAE protocols use a CNS-derived myelin peptide (e.g., MOG_35-55_) to induce an adaptive immune response against CNS tissue. However, many EAE phenotypes predominantly affect the spinal cord with relative sparing of cerebral tissue^34^. In contrast, dietary CPZ intoxication induces oxidative stress and injury preferentially affecting brain oligodendrocytes with an accompanying innate immune response^35^. Previous protocols that combined EAE and CPZ treatment have yielded robust cerebral demyelination with evidence of both innate and adaptive immune responses, but the protocols have varied and, to our knowledge, have not previously been applied to *Abcd1*-null mice. For our model, we selected two weeks of a 0.2% CPZ diet, followed by MOG peptide immunization.

Our CPZ/EAE induction produced histologic features in *Abcd1-*null mice that were similar to, but more severe than those of wild-type mice. These similarities are reminiscent of those shared by human cALD and MS lesions, which can be difficult to distinguish during the acute stages of demyelination^36^. Over time, cALD lesions distinguish themselves with a higher severity of tissue injury and persistent lesion growth, enlarging concentrically over months to years, in contrast to MS lesions which typically self-resolve within weeks^37^. We designed our experiments to investigate the brain histology in the early stage of the disease when the motor disability reached the EAE scores of 2 and 3 on days 16-22 postimmunization. Among mice evaluated up to 10 weeks after induction, we observed persistent evidence of demyelination and microglial activation (**Supplemental** Fig. 8). This should be sufficient duration of disease activity to allow for preclinical screening of potential abortive therapies for cALD. Longer duration studies are needed to establish lesion phenotypes at even later stages in our mouse model. If necessary, additional modifications, such as repeat immunization or higher concentration of pertussis, could facilitate lesion growth that is persistent but not fatal.

In humans, both cALD and MS lesions have a predilection for the medial corpus callosum^38^. The mechanisms underlying this predilection remain unknown but may be related to the region’s high myelin density, low vascularity, or other metabolic vulnerabilities^6,39,40^. Notably, the lesions induced by CPZ intoxication have a similar predilection for the medial corpus callosum in both *Abcd1^y/-^* and wild-type mice, suggesting the possibility of overlapping mechanisms that may lend value to its use in cALD and MS mouse models^35^.

The molecular mechanisms by which *ABCD1*-deficiency and its resulting VLCFA accumulation lead to the severe inflammatory demyelination in cALD remain unresolved. Inflammasomes are a family of intracellular multiprotein complexes that mediate innate immune responses and have been implicated in the pathogenesis of a growing number of neurodegenerative and metabolic diseases^41,42^. The NLRP3 inflammasome and its downstream cytokines, IL-1β and IL-18, mediate a programmed form of cell death known as pyroptosis. The NLRP3 inflammasome can be activated by many of the metabolic disruptions described in ALD, including oxidative stress, mitochondrial dysfunction, lysosomal dysfunction, and altered lipid metabolism^25,43,44^. We hypothesized that the NLRP3 inflammasome, triggered by one or more of these metabolic perturbations in ALD, might be a driver of the robust neuroinflammatory cascade in cALD. Our findings in human cALD and our cALD mouse model support our hypothesis and suggest that our mouse model may offer a preclinical tool for investigating NLRP3 as a therapeutic target for cALD.

BBB disruption and perivascular immune cell infiltration is a defining feature of cALD and is well-represented in our mouse model^6,7,39^. When the BBB is disrupted, blood-borne fibrinogen extravasates into brain parenchyma where it is converted into insoluble fibrin, a proinflammatory matrix that activates innate immune responses^45,46^. In *vitro* and animal studies demonstrated that the presence of fibrin in the brain can trigger the activation of Nlrp3/IL-18 in macrophage/microglia. The NLRP3 activation leads to inflammation and impeding the process of remyelination and neurogenesis^28,45,47^. Perivascular macrophages/microglia in our cALD mice express IL-18 at high levels. Activation of the IL-18 receptor in astrocytes converts them to the A1 phenotype, a toxic form of astrocyte that can inhibit neurite outgrowth and neurogenesis^48^. The interaction between macrophage/microglia and astrocytes through IL-18 serves as a mechanism for propagating inflammation and exacerbating neurodegenerative processes in the brain. Our model could be used for preclinical screening of therapies targeting BBB disruption, fibrin deposition, and/or the NLRP3 inflammasome.

Our cALD model may offer an opportunity to study lesion arrest and study brain remyelination. Radiographically, active human cALD brain lesions are demarcated by the presence of contrast enhancement, which indicates BBB disruption and predicts further lesion growth^6,16,38^. Our cALD mouse model recapitulates active demyelinating lesions with the presence of contrast enhancement on MRI. In human cALD, contrast enhancement and lesion growth can arrest in some patients, most often after hematopoietic stem cell transplantation. Unfortunately, the white matter injury and demyelination are irreversible^49^. In mice intoxicated with CPZ alone, acute demyelination is followed by a spontaneous remyelination phase, which typically occurs after 4-5 weeks of CPZ intoxication^35^. Our cALD mice evaluated had persistent T2 hyperintensities and contrast enhancement on MRI at 5 and 10 weeks following the initiation of a 2-week CPZ diet suggesting relatively sustained lesion activity similar to human cALD.

The current study and our proposed cALD model, in general, have important limitations. First, the actual triggers for cALD lesion formation in humans are probably varied and may involve biologically distinct pathways that are not reflected in our current induction protocol or in mouse biology more generally. Second, the current work focuses primarily on the acute phase of lesion formation, which demonstrates promising parallels to human cALD lesions. Future studies evaluating the later phases of lesion evolution will be useful in assessing the persistence of demyelination and the evolution of tissue injury. Finally, it is not yet clear from which of our proposed therapeutic targets (e.g., NLRP3, IL-18, fibrin, oxidative stress, etc.) are most promising. The current work demonstrates a causal role for Abcd1-deficiency in lesion evolution, but the causal, downstream mediators (and thus therapeutic targets) remain undefined.

## CONCLUSION

By combining *Abcd1* deficiency with CPZ diet and EAE induction, we have devised a novel mouse model of cALD that recapitulates key features of human cALD, including robust levels of cerebral demyelination, blood brain barrier disruption, perivascular infiltration of innate and adaptive immune cells, and elevated markers of oxidative stress. We also provide evidence of fibrin deposition and NLRP3 inflammasome activity in CSF and brain tissue from boys with cALD and highlight similar pathology in our cALD mouse model where fibrin deposition is robust and IL-18 expression co-localizes to perivascular macrophages/microglia. Finally, we use histological findings as well as non-invasive MRI markers to demonstrate sustained inflammation and demyelination more than two months after CPZ/EAE induction. As the first preclinical model for cALD, we expect this to enable long-sought investigations into disease mechanisms and accelerate development of candidate therapies for lesion prevention, cessation, and remyelination.

## Abbreviations

ABCD1: ATP binding cassette subfamily D member 1
cALD: Cerebral Adrenoleukodystrophy
BBB: Blood-brain barrier
CPZ: Cuprizone
EAE: Experimental Autoimmune Encephalomyelitis
LCC: Lateral Corpus Callosum,
MCC: Medial Corpus Callosum
NLRP3: NOD-, LRR- and Pyrin Domain-Containing Protein 3
PVC: Perivascular Cuff
VLCFAs: Very Long Chain Fatty Acids
X-ALD: X-linked Adrenoleukodystrophy

## Acknowledgments

We are grateful to the individuals and organizations who provided material support for this effort. This includes the generous donations of brain tissue from ALD families and from the NIH Brain and Tissue Bank. We extend our gratitude to Dr. Jay L. Degen of Cincinnati Children’s Hospital Medical Center for generously supplying us with the anti-fibrinogen antibody.

This study was supported by a training grant from NIH/NIAID to EH (5T32AR050942), the Stanford Department of Neurology, the Lucile Packard Foundation for Children’s Health, the Maternal Child Health Research Institute at Stanford, and the United Leukodystrophy Foundation ULF-2020 as well as gift funding from the Lenail-Yoler Family, the Anderson Family, the Perry Family, the Adler Family, the Taube Family, the Morgridge Family, and the Senkut Family to K.P.V, and the European Leukodystrophy Association ELA 2020-00413 to J.L.B. We are also grateful for generous funding support from the Simon Family Trust and NIH/NINDS R35 NS097976 to K.A.

## Authors contributions

EH and IS contributed equally to this work.

EH: contributed to the conception and design of the study, oversaw the generation, acquisition, analysis, and interpretation of the mouse data, wrote the initial draft of the manuscript, and approved the final version of the manuscript.

IS: contributed to the conception and design of the study, oversaw the generation, collection, assembly, analysis, and interpretation of the human data, wrote the initial draft of the manuscript.

AA: contributed to the generation, collection, assembly, analysis, and interpretation of the human data.

EY and EK: contributed to the generation, collection, assembly, analysis, and interpretation of the mouse data. AA, JK, KA, ST, PC, LP, PM, LS, KD, WR, OS, RC, PO, TL, HV, ML, and MH contributed to the generation, collection, assembly, analysis, and interpretation of the data.

JB: contributed to the conception and design of the study, contributed to the generation, collection, assembly, analysis, and interpretation of the data.

KPV: contributed to the conception and design of the study, supervised the generation, collection, assembly, analysis, and interpretation of all data, contributed to the original draft.

All authors read and approved the final manuscript.

## Potential Conflict of Interest

K.V. serves as a scientific consultant for Poxel, bluebirdbio, and Viking Therapeutics.

K.A. is the scientific founder, advisor, and shareholder of Therini Bio, Inc. Her interests are managed by Gladstone Institutes according to its conflict-of-interest policy.

## Data availability

Data presented in the manuscript is available in the article and Supplemental materials. Additional data is available from the corresponding author, KPV, upon reasonable request.

