## Supplemental Files for "A novel mouse model of cerebral adrenoleukodystrophy highlights NLRP3 activity in lesion pathogenesis"

**Annals of Neurology**

Ezzat Hashemi et al.

The supplementary includes:

Figures S1 to S9

Tables S1 to S3

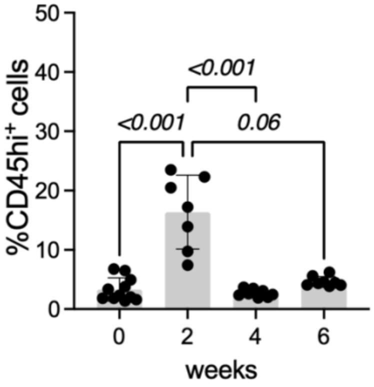


**Figure S1 Immune cell infiltration in the mouse brain at different time points of the CPZ diet.** The graph displays the ratio of CD45$^{hi}$ cells to the total immune cell population in the brain of wild-type mice using flow cytometry. The evaluation of immune cell infiltration was performed after 0, 2, 4, and 6 weeks of cuprizone diet. Bar graph illustrating the mean ± SD. One-way ANOVA and multiple comparisons determined the significance between the groups, $P<0.05$ was considered a significant difference.

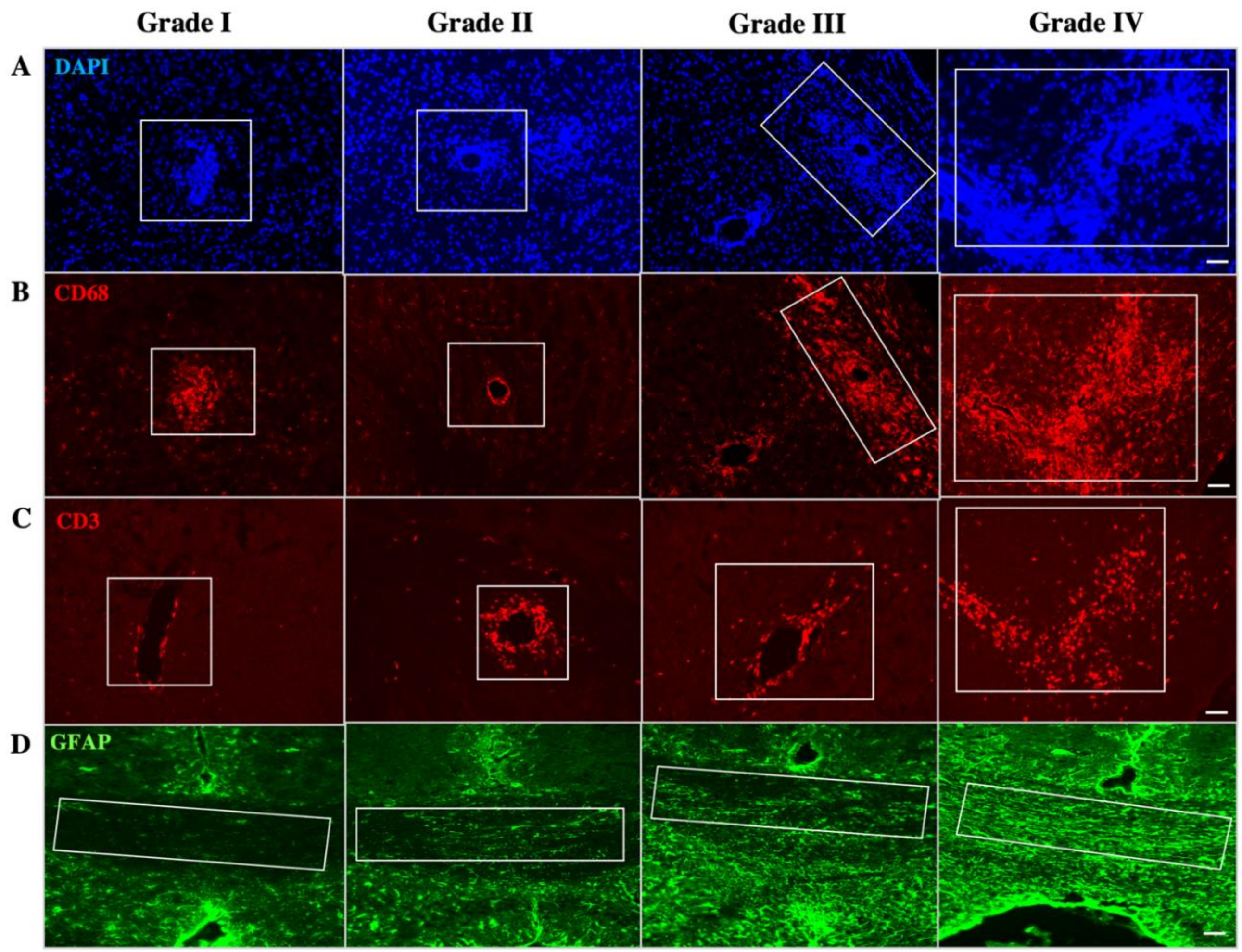


**Figure S2 Grading procedure to evaluate severity of immune cell infiltration and astrocytosis.** The severity of perivascular infiltration is graded based on the extent of immune cell penetration into the parenchyma and the lesion size. (**A**) The images display the grading of immune cell infiltration by DAPI staining. Immune cell aggregation, referred as Foci, is categorized as grade I. Immune cell trafficking around the blood vessel is assigned grade II. Grade III is assigned when immune cells penetrate the parenchyma. High infiltration and presence in more significant lesions indicate grade IV. (**B**) The images depict the grading of macrophage/microglia infiltration using CD68 staining which is comparable to DAPI staining. (**C**) The images display T cell grading. The attachment of a few T cells to the vessels without penetration, indicated as grade I. T cells accumulating around the vessels and beginning to penetrate the parenchyma are identified as grade II and III, respectively. High infiltration and T cell accumulation in large parenchyma lesion are graded as IV. The white squares represent the grade and severity of perivascular macrophage/microglia and T cells infiltration. (**D**) Astrogliosis characterized by increases in the number and/or arborization of astrocytes. The highest number and/or arborization of astrocytes in the MCC is graded as IV. The white square represents the MCC region that was graded. The scale bar represents 50 µm. MCC; Medial Corpus Callosum.

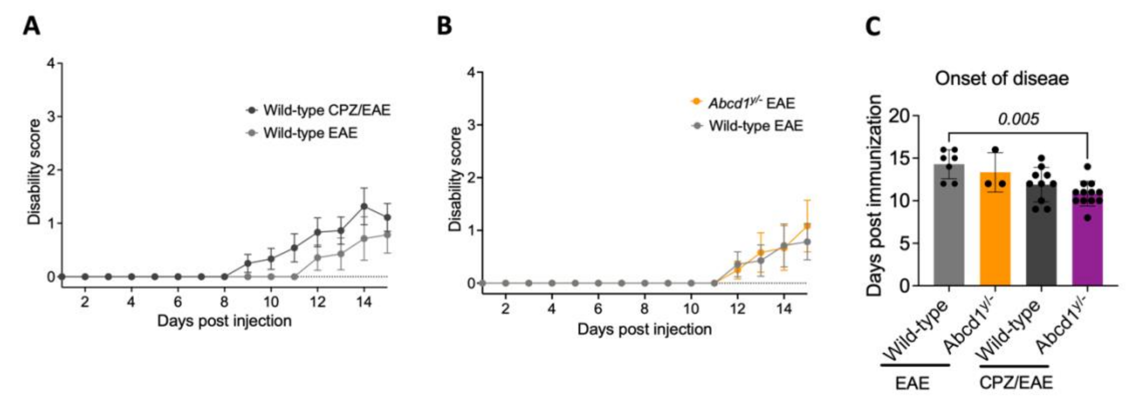


**Figure S3 Disability scores in EAE and CPZ/EAE models**.
(**A**) The disability scores in wild-type mice subjected to EAE alone and combined CPZ/EAE treatment. (**B**) The disability scores in *Abcd1*$^{y/-}$ and wild-type mice following EAE induction. (**C**) The graph displays the onset of clinical EAE scores in *Abcd1*$^{y/-}$ and wild-type mice subjected to EAE only and CPZ/EAE treatment. The bar graph represents mean ± SEM of disability scores. A significant difference between the groups was determined using a one-way ANOVA and multiple comparisons, with a significance level set at $p$<0.05.

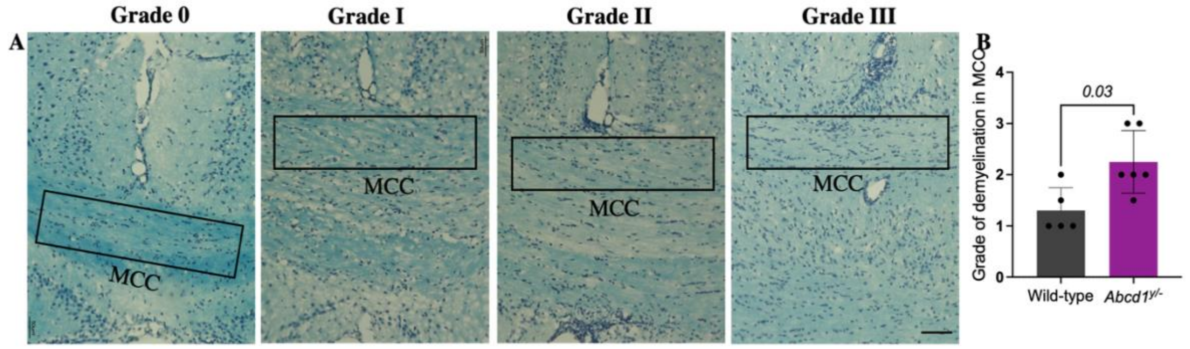


**Figure S4 Demyelination in the MCC of cALD mouse model.**
(**A**) Images indicate the grades and severity of demyelination by LFB staining. (**B**) The graph represents demyelination grades in the MCC of wild-type and *Abcd1*$^{y/-}$ mice with combined

CPZ/EAE induction (n=5-6 mice/group). The significant difference between the two groups was determined using the Mann-Whitney test, $P<0.05$ was considered a significant difference. The scale bar represents 200 µm. LFB; Luxol Fast Blue, MCC; Medial Corpus Callosum.

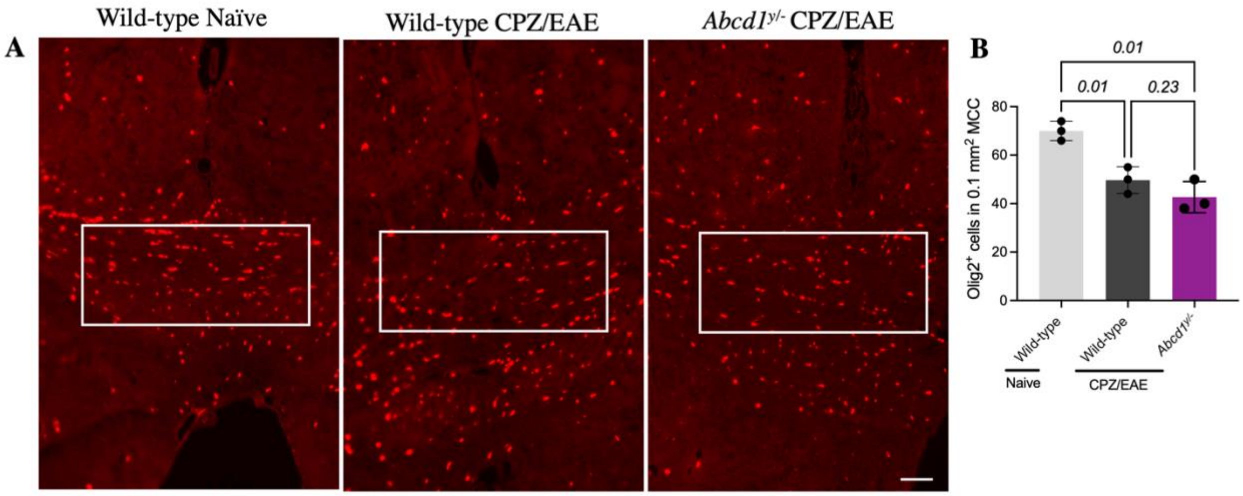


**Figure S5 Reduction of oligodendrocytes in the MCC following combined CPZ/EAE induction**. (**A**) Images indicate the presence of oligodendrocytes in the MCC of wild-type naïve mouse, as well as wild-type and *Abcd1*$^{y/-}$ mice with combined CPZ/EAE induction. (**B**) Olig2$^{+}$ cells normalized to DAPI per 0.1 mm$^2$ of MCC (n=3 mice/group). The values are presented as the mean ± SD. Statistical significance was determined using multiple comparisons and ANOVA test. $p<0.05$ was considered a significant difference. The scale bar represents 50 µm MCC; Medial Corpus Callosum.

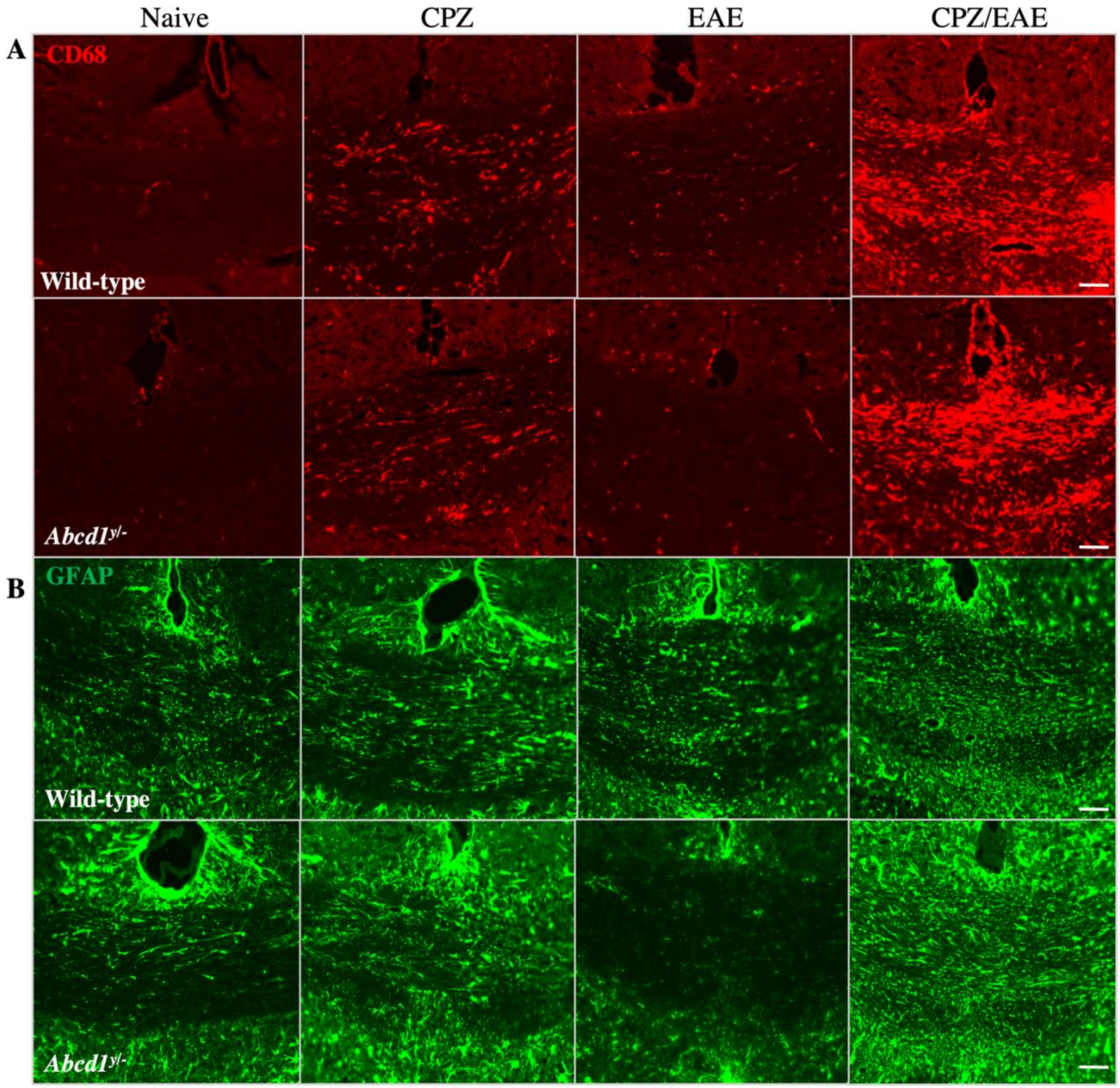


**Figure S6 Microgliosis and astrocytosis in the MCC of cALD mice**. Images display (**A**) macrophages/microglia labeled with CD68 staining and (**B**) astrocytes labeled with GFAP staining in the MCC of wild-type and *Abcd1*$^{y/-}$ mice across different conditions: naïve, CPZ, EAE, and combined CPZ/EAE induction. The scale bar represents 50 μm.

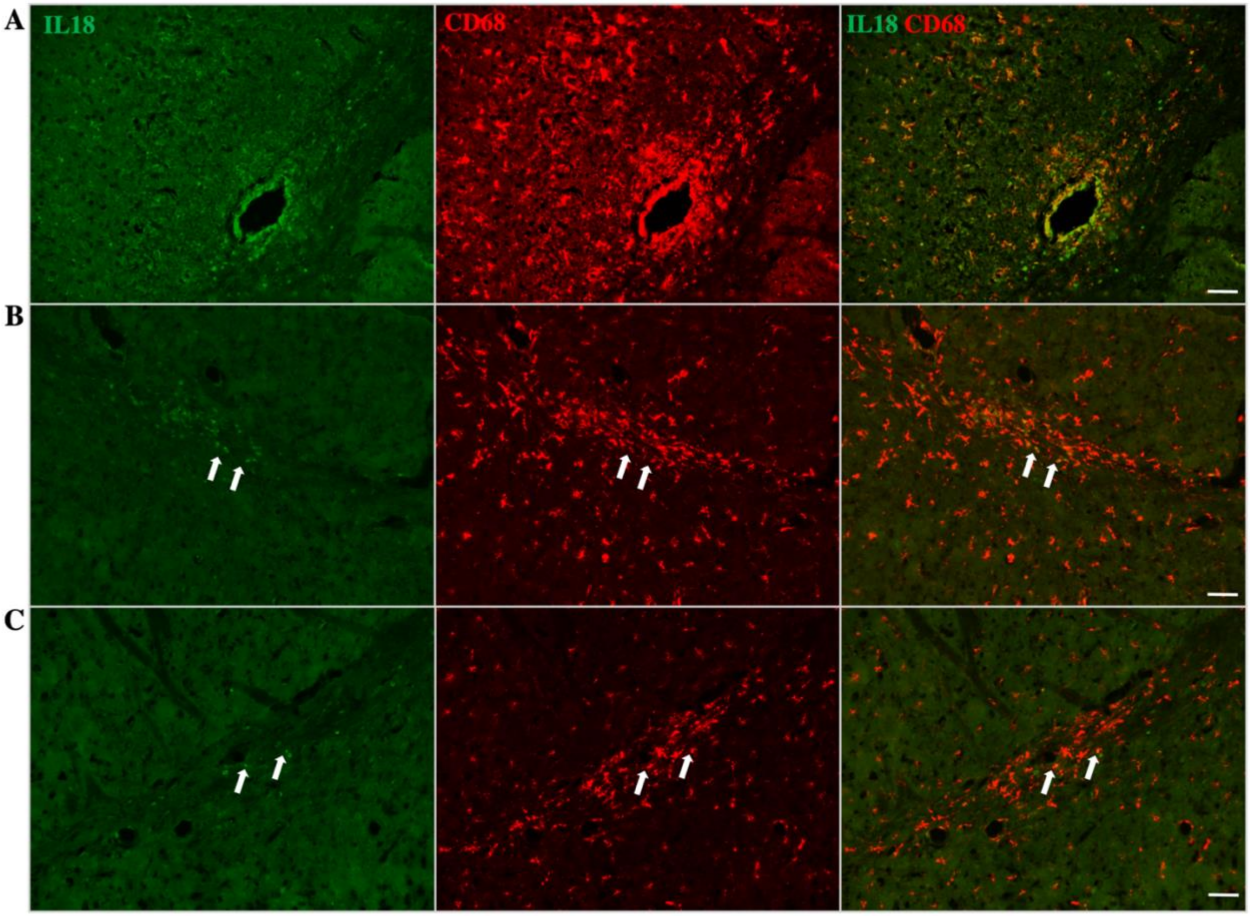


**Figure S7 Expression of IL-18 in macrophage/microglia following combined CPZ/EAE induction.** (**A**) Images depict the colocalization of IL-18 and perivascular CD68$^{+}$ cells in *Abcd1*$^{y/-}$ mouse following CPZ/EAE induction. (**B**) IL-18 expression in CD68$^{+}$ cells in the lateral corpus callosum of *Abcd1*$^{y/-}$ and (**C**) wild-type mice following CPZ/EAE induction. The scale bar represents 50 µm.

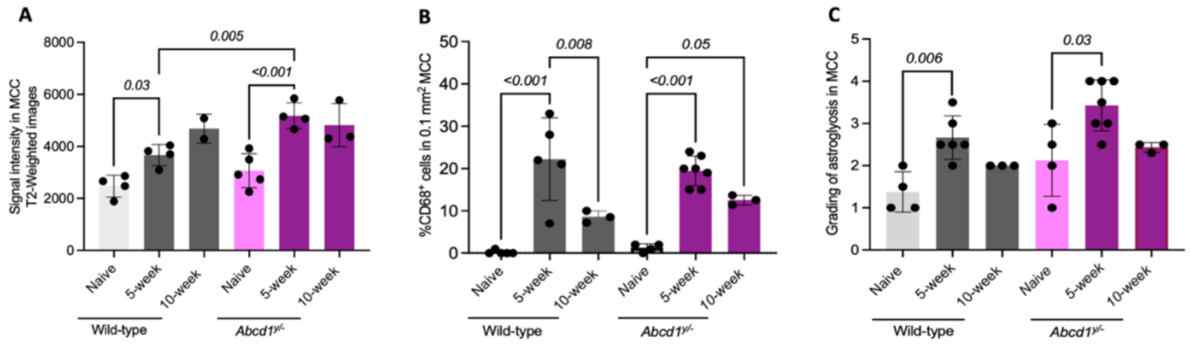


**Figure S8 Demyelination and microgliosis in later stage of combined CPZ/EAE induction.** (**A**)The graph depicts the demyelination of the MCC using signaling intensity measurements obtained from T2-weighted images. The T2 imaging was performed at three time points, including before any treatments (naïve), 5 weeks, and 10 weeks after the beginning of combined CPZ/EAE induction in both wild-type and *Abcd1*$^{y/-}$ mice. (**B**) The graph displays the percentage of macrophage/microglia cells to total cell count in the MCC of wild-type and *Abcd1*$^{y/-}$ mice, assessed through CD68 and DAPI staining. Tissue collection and histologic examinations were performed in both wild-type and *Abcd1*$^{y/-}$ mice at three specific time points: prior to any treatment (naïve), 5 weeks, and 10 weeks after initiating CPZ/EAE induction. (**C**) The graph displays astrocytosis grade in MCC using GFAP staining. The group of mice analyzed is the same as described in A section. A significant difference between the groups was determined using One-way ANOVA, and $p<0.05$ was considered statistically significant. The values are presented as the mean ± SD. MCC; Medial Corpus Callosum.

**Supplementary Table S1**: The table displays the number of mice used in each experiment across 4 arms.

| Figure | Experiments | Genotype/Treatment | Sample size |
|---|---|---|---|
| 3 | Motor disability assay | Wild-type EAE<br>*Abcd1*$^{y/-}$ EAE<br>Wild-type CPZ/EAE<br>*Abcd1*$^{y/-}$ CPZ/EAE | N=7<br>N=6<br>N=13<br>N=13 |
| 4 | T2 -weighted MRI | Wild-type naive<br>*Abcd1*$^{y/-}$ naive<br>Wild-type CPZ/EAE<br>*Abcd1*$^{y/-}$ CPZ/EAE | N=4<br>N=5<br>N=4<br>N=4 |
| 4 | MBP staining | Wild-type naive<br>*Abcd1*$^{y/-}$ naive<br>Wild-type CPZ/EAE<br>*Abcd1*$^{y/-}$ CPZ/EAE | N=4<br>N=5<br>N=5<br>N=6 |
| 4 | Fibrinogen staining | Wild-type naive<br>*Abcd1*$^{y/-}$ naive<br>Wild-type CPZ/EAE<br>*Abcd1*$^{y/-}$ CPZ/EAE | N=3<br>N=4<br>N=6<br>N=7 |
| 4 | Gp91-phox staining | Wild-type naive<br>*Abcd1*$^{y/-}$ naive<br>Wild-type CPZ/EAE<br>*Abcd1*$^{y/-}$ CPZ/EAE | N=3<br>N=4<br>N=6<br>N=7 |
| 5 | Severity of PVC | *Abcd1*$^{y/-}$ CPZ<br>*Abcd1*$^{y/-}$ EAE<br>Wild-type CPZ/EAE<br>*Abcd1*$^{y/-}$ CPZ/EAE | N=5<br>N=6<br>N=6<br>N=7 |
| 5 | Severity of perivascular CD68$^{+}$ cells | *Abcd1*$^{y/-}$ CPZ<br>*Abcd1*$^{y/-}$ EAE<br>Wild-type CPZ/EAE<br>*Abcd1*$^{y/-}$ CPZ/EAE | N=5<br>N=6<br>N=6<br>N=7 |
| 5 | Severity of perivascular CD3e$^{+}$ cells | *Abcd1*$^{y/-}$ CPZ<br>*Abcd1*$^{y/-}$ EAE<br>Wild-type CPZ/EAE<br>*Abcd1*$^{y/-}$ CPZ/EAE | N=5<br>N=6<br>N=6<br>N=7 |
| 5 | Severity of perivascular B220$^{+}$ cells | *Abcd1*$^{y/-}$ CPZ<br>*Abcd1*$^{y/-}$ EAE<br>Wild-type CPZ/EAE<br>*Abcd1*$^{y/-}$ CPZ/EAE | N=5<br>N=6<br>N=6<br>N=7 |
| 6 | Microgliosis in MCC (CD68 staining) | Wild-type naive<br>*Abcd1*$^{y/-}$ naive<br>Wild-type CPZ<br>*Abcd1*$^{y/-}$ CPZ<br>Wild-type EAE<br>*Abcd1*$^{y/-}$ EAE<br>Wild-type CPZ/EAE<br>*Abcd1*$^{y/-}$ CPZ/EAE | N=5<br>N=4<br>N=4<br>N=3<br>N=4<br>N=3<br>N=5<br>N=7 |

| | | | |
|---|---|---|---|
| 6 | Astrocytosis in MCC (GFAP staining) | Wild-type naive<br>*Abcd1*$^{y/-}$ naive<br>Wild-type CPZ<br>*Abcd1*$^{y/-}$ CPZ<br>Wild-type EAE<br>*Abcd1*$^{y/-}$ EAE<br>Wild-type CPZ/EAE<br>*Abcd1*$^{y/-}$ CPZ/EAE | N=4<br>N=4<br>N=3<br>N=3<br>N=3<br>N=5<br>N=6<br>N=7 |
| 1S | %CD45$^{hi}$ /total immune cells in brain (Flowcytometry) | Wild-type naive<br>Wild-type 2-week CPZ<br>Wild-type 4-week CPZ<br>Wild-type 6-week CPZ | N=11<br>N=7<br>N=7<br>N=8 |
| 3S | Onset of disease in EAE and CPZ/EAE treatments | Wild-type EAE<br>*Abcd1*$^{y/-}$ EAE<br>Wild-type CPZ/EAE<br>*Abcd1*$^{y/-}$ CPZ/EAE | N=7<br>N=3<br>N=10<br>N=12 |
| 4S | LFB staining | Wild-type CPZ/EAE<br>*Abcd1*$^{y/-}$ CPZ/EAE | N=5<br>N=6 |
| 5S | Olig2 staining | Wild-type naive<br>Wild-type CPZ/EAE<br>*Abcd1*$^{y/-}$ CPZ/EAE | N=3<br>N=3<br>N=3 |
| 8S | T2-weighted MRI | Wild-type naive<br>Wild-type CPZ/EAE 5-week<br>Wild-type CPZ/EAE 10-week<br>*Abcd1*$^{y/-}$ naive<br>*Abcd1*$^{y/-}$ CPZ/EAE 5-week<br>*Abcd1*$^{y/-}$ CPZ/EAE 10-week | N=4<br>N=4<br>N=2<br>N=5<br>N=4<br>N=3 |
| 8S | Microgliosis in MCC (CD68 staining) | Wild-type naive<br>Wild-type CPZ/EAE 5-week<br>Wild-type CPZ/EAE 10-week<br>*Abcd1*$^{y/-}$ naive<br>*Abcd1*$^{y/-}$ CPZ/EAE 5-week<br>*Abcd1*$^{y/-}$ CPZ/EAE 10-week | N=5<br>N=5<br>N=3<br>N=4<br>N=7<br>N=3 |
| 8S | Astrocytosis in MCC (GFAP staining) | Wild-type naive<br>Wild-type CPZ/EAE 5-week<br>Wild-type CPZ/EAE 10-week<br>*Abcd1*$^{y/-}$ naive<br>*Abcd1*$^{y/-}$ CPZ/EAE 5-week<br>*Abcd1*$^{y/-}$ CPZ/EAE 10-week | N=4<br>N=6<br>N=3<br>N=4<br>N=7<br>N=3 |

**Supplementary Table S2.** The table lists information about the primary and secondary antibodies used for immunohistochemistry.

| **Antibody** | **Supplier** | **Host** | **Dilution** | **Cat. Number** |
|---|---|---|---|---|
| CD3e | BD Biosciences | Hamster | 1:300 | 550277 |
| CD45R/B220 | BD Biosciences | Rat | 1:300 | 553085 |
| CD68 | Bio-Rad | Rat | 1:300 | MCA1957GA |
| GFAP | DAKO | Rabbit | 1:600 | Z0334 |
| GP91-phox | BD Bioscience | Mouse | 1:200 | 611414 |
| Iba1 | Wako | Rabbit | 1:100 | 019-19741 |
| IL-18 | Protein Tech | Mouse | 1:200 | 60070-1-Ig |
| IL-18 | Abcam | Rabbit | 1:100 | ab191152 |
| IL-18 | Invitrogen | Rabbit | 1/100 | PA5-79481 |
| MBP | Abcam | Rat | 1:300 | AB7349 |
| OLIG2 | Millipore | Mouse | 1:400 | MABN50 |
| PLP | Abcam | Rabbit | 1:300 | AB28486 |
| Alexa Flour™ 555 | Invitrogen | Goat anti-mouse | 1:1000 | A21424 |
| Alexa Flour™ 555 | Invitrogen | Goat anti-rat | 1:1000 | A21434 |
| Alexa Flour™ 555 | Invitrogen | Goat anti-rabbit | 1:1000 | A21428 |
| Alexa Flour™ 647 | Invitrogen | Goat anti-rabbit | 1:1000 | A27040 |
| Alexa Flour™ 647 | Sigma-Aldrich | Goat anti-mouse | 1:1000 | A32728 |
| Alexa Flour™ 647 | Invitrogen | Goat anti-hamster | 1:1000 | A21451 |
| Alexa Flour™ 488 | Invitrogen | Goat anti-rabbit | 1:1000 | A-11008 |
| Streptoavidin 488 | Invitrogen | Not Applicable | 1/700 | S32354 |

**Supplementary Table S3**. The table indicates the homogeneity of variances and mean ± SD for each immunostaining and T2-weighted MRI test.

| Figure | Experiments | Homogeneity of Variances | Genotype/Treatment | Mean ± SD |
|---|---|---|---|---|
| 4 | T2 -weighted MRI | Brown-Forsythe and Welch ANOVA | Wild-type naive<br>*Abcd1*$^{y/-}$ naive<br>Wild-type CPZ/EAE<br>*Abcd1*$^{y/-}$ CPZ/EAE | 2474 ± 421<br>3086 ± 655<br>3667 ± 407<br>5185 ± 493 |
| 4 | MBP staining | Ordinary One-Way ANOVA | Wild-type naive<br>*Abcd1*$^{y/-}$ naive<br>Wild-type CPZ/EAE<br>*Abcd1*$^{y/-}$ CPZ/EAE | 78.75 ± 4.11<br>75.6 ± 6.22<br>59.38 ± 4.80<br>50.42 ± 7.21 |
| 4 | Fibrinogen staining | Brown-Forsythe and Welch ANOVA | Wild-type naive<br>*Abcd1*$^{y/-}$ naive<br>Wild-type CPZ/EAE<br>*Abcd1*$^{y/-}$ CPZ/EAE | 0.10 ± 0.09<br>0.11 ± 0.08<br>0.91 ± 0.62<br>2.4 ± 0.84 |
| 4 | Gp91-phox staining | Brown-Forsythe and Welch ANOVA | Wild-type naive<br>*Abcd1*$^{y/-}$ naive<br>Wild-type CPZ/EAE<br>*Abcd1*$^{y/-}$ CPZ/EAE | 0.03 ± 0.02<br>0.02 ± 0.01<br>0.82 ± 0.60<br>1.77 ± 0.26 |
| 5 | Severity of PVC | Brown-Forsythe and Welch ANOVA | *Abcd1*$^{y/-}$ CPZ<br>*Abcd1*$^{y/-}$ EAE<br>Wild-type CPZ/EAE<br>*Abcd1*$^{y/-}$ CPZ/EAE | 0.07 ± 0.08<br>0.05 ± 0.05<br>0.75 ± 0.60<br>2.21 ± 0.74 |
| 5 | Severity of perivascular CD68$^{+}$ cells | Brown-Forsythe and Welch ANOVA | *Abcd1*$^{y/-}$ CPZ<br>*Abcd1*$^{y/-}$ EAE<br>Wild-type CPZ/EAE<br>*Abcd1*$^{y/-}$ CPZ/EAE | 0.05 ± 0.05<br>0.07 ± 0.08<br>0.48 ± 0.45<br>2.11 ± 0.99 |
| 5 | Severity of perivascular CD3e$^{+}$ cells | Unpaired t-test | Wild-type CPZ/EAE<br>*Abcd1*$^{y/-}$ CPZ/EAE | 0.45 ± 0.43<br>1.6 ± 0.62 |
| 5 | Severity of perivascular B220$^{+}$ cells | Mann-Whitney test | Wild-type CPZ/EAE<br>*Abcd1*$^{y/-}$ CPZ/EAE | 0.18 ± 0.21<br>1.1± 0.51 |
| 6 | Microgliosis in MCC (CD68 staining) | Brown-Forsythe and Welch ANOVA | Wild-type CPZ<br>*Abcd1*$^{y/-}$ CPZ<br>Wild-type EAE<br>*Abcd1*$^{y/-}$ EAE<br>Wild-type CPZ/EAE<br>*Abcd1*$^{y/-}$ CPZ/EAE | 15.3 ± 6.34<br>16.3 ± 3.51<br>3.50 ± 2.08<br>3.0 ± 1.0<br>22.2 ± 9.78<br>19.4 ± 3.51 |
| 6 | Astrocytosis in MCC (GFAP staining) | Brown-Forsythe and Welch ANOVA | Wild-type naive<br>*Abcd1*$^{y/-}$ naive<br>Wild-type CPZ<br>*Abcd1*$^{y/-}$ CPZ<br>Wild-type EAE<br>*Abcd1*$^{y/-}$ EAE<br>Wild-type CPZ/EAE | 1.38 ± 0.48<br>2.13 ± 0.85<br>3.5 ± 0.5<br>3 ± 0.5<br>2.33 ± 0.76<br>1.38 ± 0.48 |

| | | | | |
|---|---|---|---|---|
| | | | *Abcd1*$^{y/-}$ CPZ/EAE | 2.7 ± 0.52<br>3.42 ± 0.61 |
| 1S | %CD45$^{hi}$ /total immune cells in brain, Flowcytometry | Kruskal-Wallis test | Wild-type naive<br>Wild-type 2-week CPZ<br>Wild-type 4-week CPZ<br>Wild-type 6-week CPZ | 3.31 ± 1.96<br>16.4 ± 6.24<br>2.70 ± 0.66<br>4.64 ± 0.83 |
| 3S | Onset of disease in EAE and CPZ/EAE treatments | Kruskal-Wallis test | Wild-type EAE<br>*Abcd1*$^{y/-}$ EAE<br>Wild-type CPZ/EAE<br>*Abcd1*$^{y/-}$ CPZ/EAE | 14.3 ± 1.7<br>13.3 ± 2.31<br>11.9 ± 2.02<br>10.8 ± 1.47 |
| 4S | LFB staining | Mann-Whitney test | Wild-type CPZ/EAE<br>*Abcd1*$^{y/-}$ CPZ/EAE | 1.0 ± 0.04<br>2.0 ± 0.6 |
| 5S | Olig2 staining | Brown-Forsythe and Welch ANOVA | Wild-type naive<br>Wild-type CPZ/EAE<br>*Abcd1*$^{y/-}$ CPZ/EAE | 70 ± 4.0<br>50 ± 5.5<br>43 ± 6.4 |
| 8S | T2-weighted MRI | Ordinary One-Way ANOVA | Wild-type naive<br>Wild-type CPZ/EAE 5-week<br>*Abcd1*$^{y/-}$ naive<br>*Abcd1*$^{y/-}$ CPZ/EAE 5-week | 2474 ± 421<br>3667 ± 407<br>3068 ± 655<br>5185 ± 493 |
| 8S | Microgliosis in MCC (CD68 staining) | Ordinary One-Way ANOVA | Wild-type naive<br>Wild-type CPZ/EAE 5-week<br>Wild-type CPZ/EAE 10-week<br>*Abcd1*$^{y/-}$ naive<br>*Abcd1*$^{y/-}$ CPZ/EAE 5-week<br>*Abcd1*$^{y/-}$ CPZ/EAE 10-week | 0.2 ± 0.45<br>22.2 ± 9.78<br>8.57 ± 1.4<br>1.25 ± 0.96<br>19.4 ± 3.51<br>12.5 ± 1.13 |
| 8S | Astrocytosis in MCC (GFAP staining) | Kruskal-Wallis test | Wild-type naive<br>Wild-type CPZ/EAE 5-week<br>*Abcd1*$^{y/-}$ naive<br>*Abcd1*$^{y/-}$ CPZ/EAE 5-week | 1.38 ± 0.48<br>2.67 ± 0.52<br>2.13 ± 0.85<br>3.43 ± 0.61 |

| Authors | Contribution | Affiliation |
| --- | --- | --- |
| Ezzat Hashemi, PhD | contributed to the conception and design of the study, oversaw the generation, acquisition, analysis, and interpretation of the mouse data, wrote the initial draft of the manuscript, and approved the final version of the manuscript. | Department of Neurology and Neurological Sciences, Stanford University School of Medicine, Stanford, CA, USA |
| Isha Narain Srivastava, MD, PhD | contributed to the conception and design of the study, oversaw the generation, collection, assembly, analysis, and interpretation of the human data, wrote the initial draft of the manuscript. | Department of Neurology and Neurological Sciences, Stanford University School of Medicine, Stanford, CA, USA |
| Alejandro Aguirre MD | contributed to the generation, collection, assembly, analysis, and interpretation of the human data. | Department of Neurology and Neurological Sciences, Stanford University School of Medicine, Stanford, CA, USA |
| Ezra Tilahan Yoseph, BS | contributed to the generation, collection, assembly, analysis, and interpretation of the mouse data. | Department of Neurology and Neurological Sciences, Stanford University School of Medicine, Stanford, CA, USA |
| Esha Kaushal PhD | contributed to the generation, collection, assembly, analysis, and interpretation of the mouse data. | Department of Neurology and Neurological Sciences, Stanford University School of Medicine, Stanford, CA, USA |
| Avni Awani, PhD | contributed to the generation, collection, assembly, analysis, and interpretation of the data. | Department of Neurology and Neurological Sciences, Stanford University School of Medicine, Stanford, CA, USA |
| Jae Kyu. Ryu, PhD | contributed to the generation, collection, assembly, analysis, and interpretation of the data. | Gladstone Institute for Neurological Disease; San Francisco, CA, USA. Center for Neurovascular Brain Immunology at Gladstone and UCSF; San Francisco, CA USA. |

| | | |
|---|---|---|
| | | Department of Neurology, Weill Institute for Neurosciences, University of California San Francisco; San Francisco, CA, USA. |
| Katerina Akassoglou, PhD | contributed to the generation, collection, assembly, analysis, and interpretation of the data. | Gladstone Institute for Neurological Disease; San Francisco, CA, USA. Center for Neurovascular Brain Immunology at Gladstone and UCSF; San Francisco, CA USA. Department of Neurology, Weill Institute for Neurosciences, University of California San Francisco; San Francisco, CA, USA. |
| Shahrzad Talebian M.Sc | contributed to the generation, collection, assembly, analysis, and interpretation of the data. | Department of Neurology and Neurological Sciences, Stanford University School of Medicine, Stanford, CA, USA. |
| Pauline Chu B.S, HT | contributed to the generation, collection, assembly, analysis, and interpretation of the data. | Stanford Human Research Histology Core, Stanford University School of Medicine, Stanford, CA, USA. |
| Laura Pisani PhD | contributed to the generation, collection, assembly, analysis, and interpretation of the data. | Department of Radiology, Stanford University School of Medicine Stanford, CA, USA. |
| Patricia Musolino MD, PhD | contributed to the generation, collection, assembly, analysis, and interpretation of the data. | Department of Neurology, Massachusetts General Hospital, Boston, MA, USA. |

| | | |
|---|---|---|
| | | Center for Genomic Medicine, Massachusetts General Hospital, Boston, MA, USA. |
| Lawrence Steinman, MD | contributed to the generation, collection, assembly, analysis, and interpretation of the data. | Department of Neurology and Neurological Sciences, Stanford University School of Medicine, Stanford, CA, USA. |
| Kristian Doyle, PhD | contributed to the generation, collection, assembly, analysis, and interpretation of the data. | Department of Immunobiology, University of Arizona, Tucson, AZ, USA. |
| William H Robinson MD, PhD | contributed to the generation, collection, assembly, analysis, and interpretation of the data. | Department of Immunology & Rheumatology, Stanford University School of Medicine, Stanford, CA, USA. |
| Orr Sharpe, MSc | contributed to the generation, collection, assembly, analysis, and interpretation of the data. | Department of Immunology & Rheumatology, Stanford University School of Medicine, Stanford, CA, USA. |
| Romain Cayrol, MD, PhD | contributed to the generation, collection, assembly, analysis, and interpretation of the data. | Department of Pathology, Clinical Department of Laboratory Medicine, University of Montreal, Quebec, Canada. |
| Paul Orchard, MD | contributed to the generation, collection, assembly, analysis, and interpretation of the data. | Division of Pediatric Blood & Marrow Transplantation, University of Minnesota, Minneapolis, MN, USA. |

| | | |
|---|---|---|
| Troy Lund, MD, PhD | contributed to the generation, collection, assembly, analysis, and interpretation of the data. | Division of Pediatric Blood & Marrow Transplantation, University of Minnesota, Minneapolis, MN, USA. |
| Hannes Vogel, MD | contributed to the generation, collection, assembly, analysis, and interpretation of the data. | Departments of Pathology, Stanford University School of Medicine, Stanford, CA, USA. |
| Max Lenail, BS | | |
| May Htwe Han MD | contributed to the generation, collection, assembly, analysis, and interpretation of the data. | Department of Neurology and Neurological Sciences, Stanford University School of Medicine, Stanford, CA, USA |
| Joshua Leith Bonkowsky MD, PhD | contributed to the conception and design of the study, contributed to the generation, collection, assembly, analysis, and interpretation of the data. | Division of Pediatric Neurology, Department of Pediatrics, University of Utah School of Medicine, Salt Lake City, Utah.<br>Brain and Spine Center, Primary Children's Hospital, Salt Lake City, Utah.<br>Primary Children's Center for Personalized Medicine, Salt Lake City, Utah |
| Keith P. Van Haren, MD | contributed to the conception and design of the study, supervised the generation, collection, assembly, analysis, and interpretation of all data, contributed to the original draft.<br>All authors read and approved the final manuscript. | Department of Neurology and Neurological Sciences, Stanford University School of Medicine, Stanford, CA, USA.<br>Department of Pediatrics, Stanford University School of Medicine, Stanford, CA, USA |
